# Integrated DNA methylation and gene expression profiling across multiple brain regions implicate novel genes in Alzheimer’s disease

**DOI:** 10.1101/430603

**Authors:** Stephen A. Semick, Rahul A. Bharadwaj, Leonardo Collado-Torres, Ran Tao, Joo Heon Shin, Amy Deep-Soboslay, James Weiss, Daniel R. Weinberger, Thomas M. Hyde, Joel E. Kleinman, Andrew E. Jaffe, Venkata S. Mattay

## Abstract

**Background:** Late-onset Alzheimer’s disease (AD) is a complex age-related neurodegenerative disorder that likely involves epigenetic factors. To better understand the epigenetic state associated with AD represented as variation in DNA methylation (DNAm), we surveyed 420,852 DNAm sites from neurotypical controls (N=49) and late-onset AD patients (N=24) across four brain regions (hippocampus, entorhinal cortex, dorsolateral prefrontal cortex and cerebellum).

**Results:** We identified 858 sites with robust differential methylation, collectively annotated to 772 possible genes (FDR<5%, within 10kb). These sites were overrepresented in AD genetic risk loci (p=0.00655), and nearby genes were enriched for processes related to cell-adhesion, immunity, and calcium homeostasis (FDR<5%). We analyzed corresponding RNA-seq data to prioritize 130 genes within 10kb of the differentially methylated sites, which were differentially expressed and had expression levels associated with nearby DNAm levels (p<0.05). This validated gene set includes previously reported (e.g. *ANK1, DUSP22*) and novel genes involved in Alzheimer’s disease, such as *ANKRD30B*.

**Conclusions:** These results highlight DNAm changes in Alzheimer’s disease that have gene expression correlates, implicating DNAm as an epigenetic mechanism underlying pathological molecular changes associated with AD. Furthermore, our framework illustrates the value of integrating epigenetic and transcriptomic data for understanding complex disease.

## Introduction

Alzheimer’s disease (AD) is an age-related neurodegenerative disease of complex etiology and is characterized by a progressive decline of memory and cognitive faculties. Although late-onset AD risk has a strong genetic component including the APOE locus [6] and several other loci identified through genome-wide association studies (GWAS) [24], disease risk is also influenced by lifestyle and environmental factors such as diet [48], sleep [26], education and literacy [22], history of head trauma [22], and level of physical activity [41]. Epigenetic marks such as DNA methylation (DNAm) can capture molecular regulatory mechanisms through which environmental and lifestyle factors may modulate AD risk, including through interaction with underlying genetic risk. A better understanding of the epigenetic regulation of gene expression in late-onset Alzheimer’s disease (LOAD) could facilitate the discovery of more viable preventive and therapeutic strategies for this devastating disease.

While previous studies have identified global DNA methylation alterations in Alzheimer’s disease [4, 5, 34], recent landmark epigenome-wide association studies (EWAS) [2, 8, 31] have identified DNAm changes around specific genes. In a study using post-mortem brain tissue from the dorsolateral prefrontal cortex (DLPFC), 71 genome-wide sites had DNAm levels that robustly associated with AD neuropathology, ultimately implicating seven genes within 50kb with dysregulated expression: *CDH23, DIP2A, RHBDF2, RPL13, SERPINF1, SERPINF2*, and *ANK1* [8]. In a parallel study, DNAm of a region near *ANK1* strongly correlated with neuropathological measures in three cortical brain regions (entorhinal cortex, superior temporal gyrus, and prefrontal cortex), but not cerebellum, suggesting some epigenetic perturbations in AD occur across multiple cortical regions [31]. Likewise, a 48kb region within the *HOXA* gene cluster was differentially methylated in AD across multiple cortical regions [50]. Subsequent studies [18, 46, 49, 51, 53] have integrated DNAm and genetic evidence to highlight AD relevant genes. Thus far, however, these approaches have implicated only a few genes associated with DNAm variation in Alzheimer’s disease.

While these previous studies have successfully identified DNAm differences associated with AD, they have been limited in their ability to connect these epigenetic differences to corresponding gene expression changes. Previous analyses often used targeted qPCR for a small number of transcripts and did not comprehensively survey gene expression changes among all implicated DNAm loci. Moreover, the relationship between DNAm and gene expression has remained unclear from earlier studies because each data type originated from non-overlapping subjects. Integrating DNA methylation with an unbiased method for surveying the transcriptome—RNA sequencing (RNA-seq)— may therefore offer deeper insight into [15] epigenetically mediated transcriptional dysregulation associated with Alzheimer’s disease.

We therefore performed multi-stage analyses incorporating paired DNA methylation and gene expression data. Given the unique cellular composition and potentially distinct susceptibilities to Alzheimer’s disease neuropathology of different brain regions, we chose to study four regions from each brain donor: dorsolateral prefrontal cortex (DLPFC), entorhinal cortex (ERC), hippocampus (HIPPO) all three previously implicated in AD; and the cerebellar cortex (CRB). In our first stage, we compared the DNAm landscape of neurotypical controls to late-onset AD, with the Illumina’s Human Methylation 450k (HM450k) array. Then, for genes adjacent to DNAm loci identified from this analysis, we analyzed RNA seq data for differential expression between AD cases and controls, as well as for correlation between DNAm and gene expression levels. This undertaking represents, to our knowledge, one of the most comprehensive integrations of epigenetic and transcriptomic data for LOAD in postmortem human brain tissue to date.

## Results

### Clinical characteristics of AD subjects compared to unaffected controls

The dataset used for our epigenome-wide scan of AD consisted of 98 post-mortem brain donors with methylation data generated from four brain regions: entorhinal cortex (ERC), dorsolateral prefrontal cortex (DLPFC), hippocampus (HIPPO), and cerebellum (CRB) (**Table S1**). 30 donors had intermediate signs of pathology not meeting our criteria for Alzheimer’s disease (AD) and were used in secondary analyses (see Methods). Therefore, our discovery cohort includes 49 neurotypical controls and 24 AD donors with a neuropathological diagnosis (see Methods) as estimated by standard Braak staging and CERAD scoring. Compared to controls, AD donors were older (p=2.95×10^−8^, t_two-tailed_=7.67) and had reduced overall brain mass (p=9.33×10^−6^, t_two-tailed_ =5.24, **Figure S1**). APOE risk, defined here as the number of ε4 alleles, is a strong genetic factor underlying AD clinical risk [9, 29] and was indeed more common in our AD samples than our control samples (p=2.99×10^−5^, Fisher’s Exact Test). Alzheimer’s disease was not significantly associated with differences in DNAm-estimated NeuN+ (neuronal) composition in any of the four brain regions in our sample (p>0.05, **Results S1, Figure S2**), in line with previous large epigenome-wide association studies [8]. There was also no significant association between epigenetic age acceleration and AD in any of the brain regions (p>0.05 **Result S2, Figures S3 and S4**).

### *CpG-site* DNAm differences between AD subjects and unaffected controls

To identify differentially methylated sites with “shared effects” across multiple brain regions [31, 50], we adopted a powerful cross-brain region strategy (**Results S3**, see Methods) where we analyzed all of our samples in conjunction (N=269 total DNAm samples; **Table S1**). With this cross-region analysis, we identified 858 DNAm sites differentially methylated between AD and controls (FDR<5%, **Table S4**). Collectively, these sites were within 10kb of 1,156 Gencode-annotated genes (v25), had a small median absolute difference in DNAm of 3.98%, and were relatively more methylated in subjects with AD compared to controls (N=491, 57.2%, p=2.30×10^−5^, **Figure 1A**). This analysis identified CpGs near genes *ANK1* and *MYO1C*, genes that have been previously associated[8, 31] with AD neuropathology, as well as an association with *ANKRD30B* and other genes (**Figure 1B**). Among the most significant methylation sites, one near *WDR81* and *SERPINF2* has been previously reported (within 10kb, p=3.48×10^−11^, cg19803550, 4^th^ most significant, **Table S4**)[8]. When we hierarchically clustered adjusted beta values of these 858 significant DMPs for all of our samples (N=269), we found that samples clustered primarily by cerebellar vs. non-cerebellar brain region, then secondly by AD diagnosis (**Figure 1C**). The post-hoc region-specific statistics were highly correlated across all four brain regions at these sites, confirming our expectation that AD has similar associations with DNA methylation at many of these sites across the different brain regions (**Figure S5**).

**Figure 1A.**
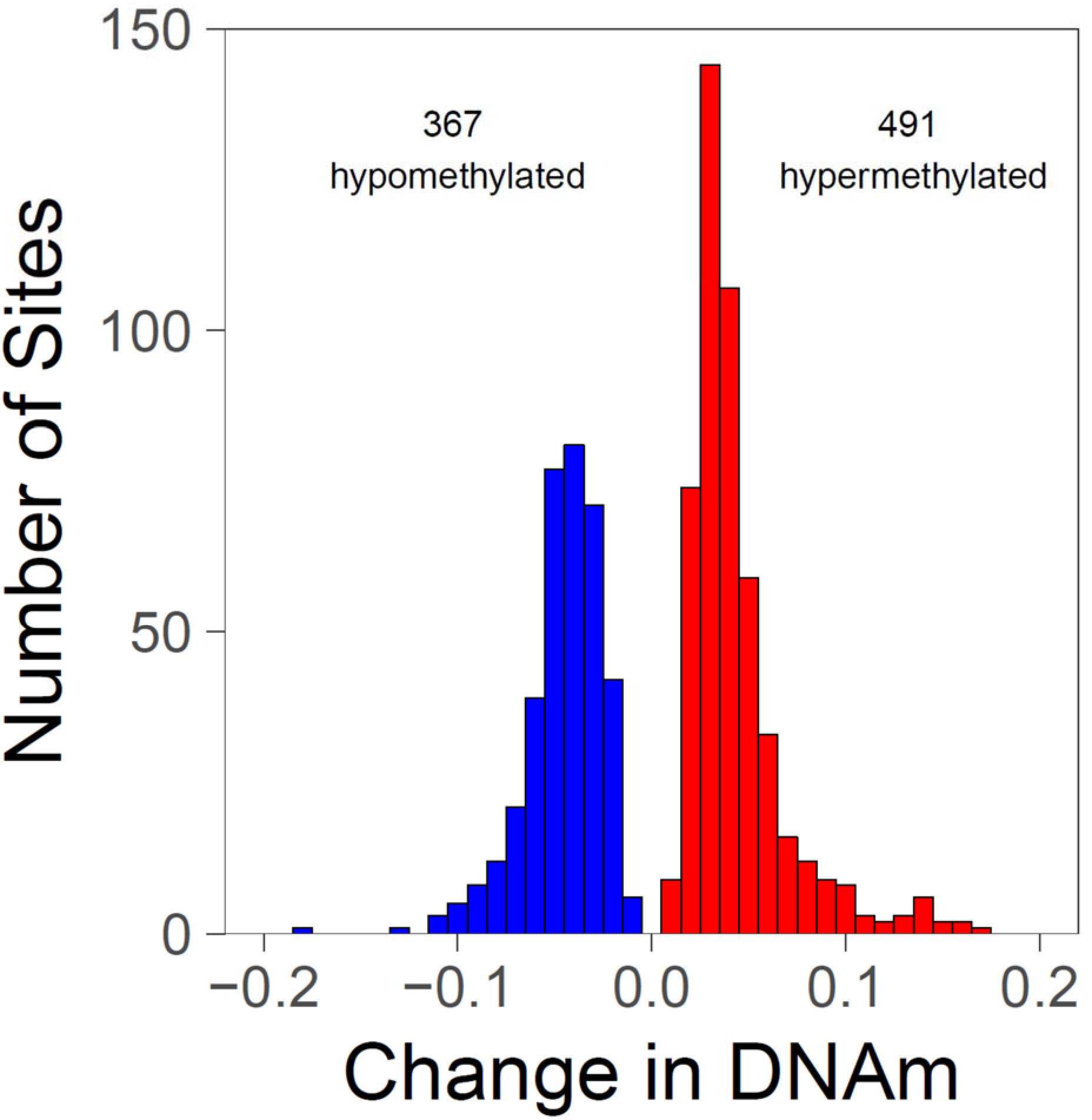
Histogram of effect sizes for significant differentially methylated probes (DMP). 858 DNAm sites are significantly differentially methylated with the cross-region model (at false discovery rate, FDR<5%). Of these 858 sites, 367 are less methylated (hypomethylated) and 491 are more methylated (hypermethylated) in AD patients compared to unaffected controls. The greater number of hypermethylated sites constitutes statistically significant enrichment (p=2.30×10^−5^).

**Figure 1B.**
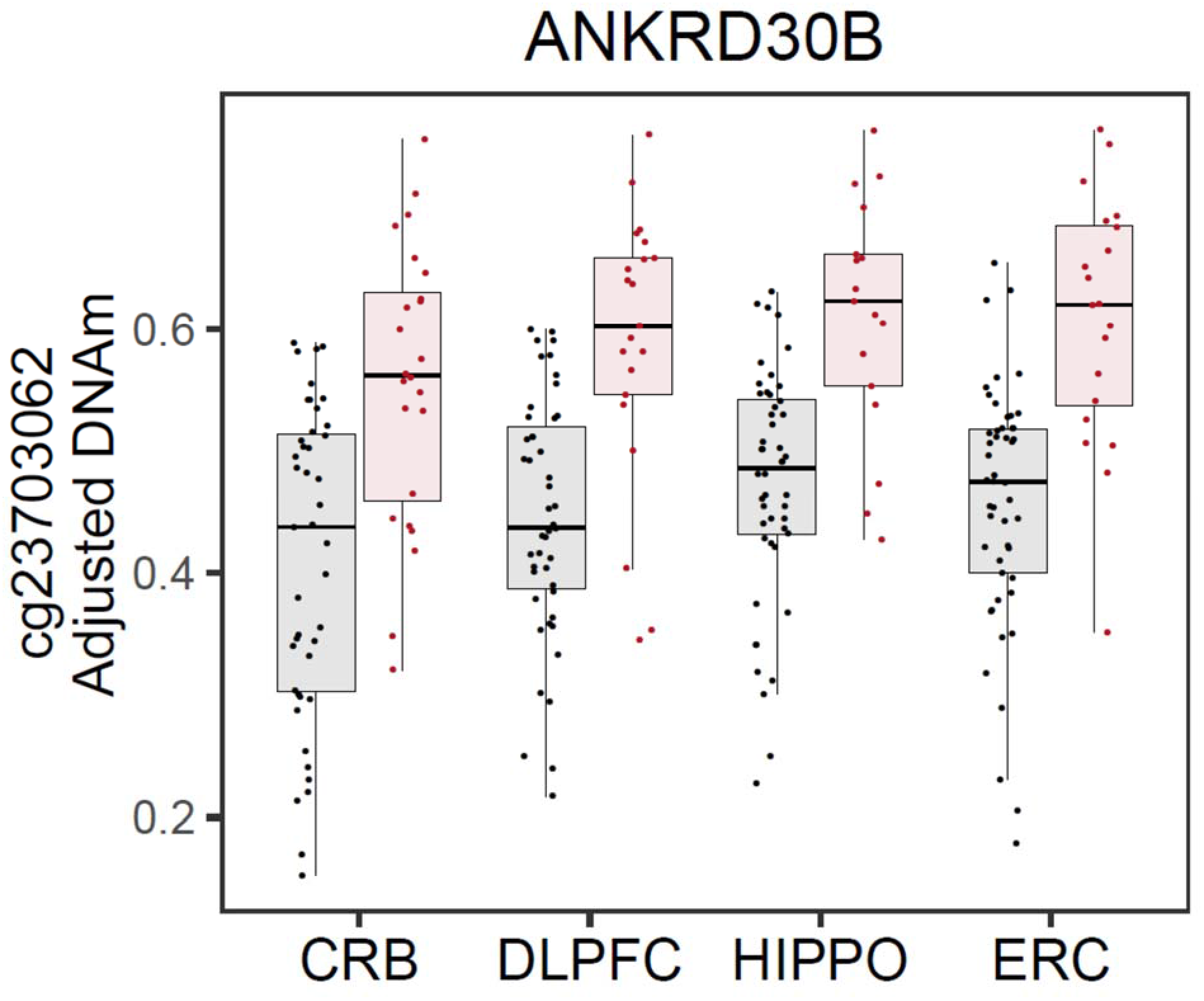
*ANKRD30B* differentially methylated probes (DMP) hypermethylation. Black points represent unaffected control samples and red points represent samples with a diagnosis of symptomatic Alzheimer’s disease (AD). Under the cross-region model, a DNAm site near *ANKRD30B* was significantly more methylated in AD samples (red) than unaffected controls (black). Plotted beta values were adjusted for age, sex, ancestry, and estimates of technical variation using a linear model with logit transformation. *ANKRD30B:* Ankyrin Repeat Domain 30B

**Figure 1C.**
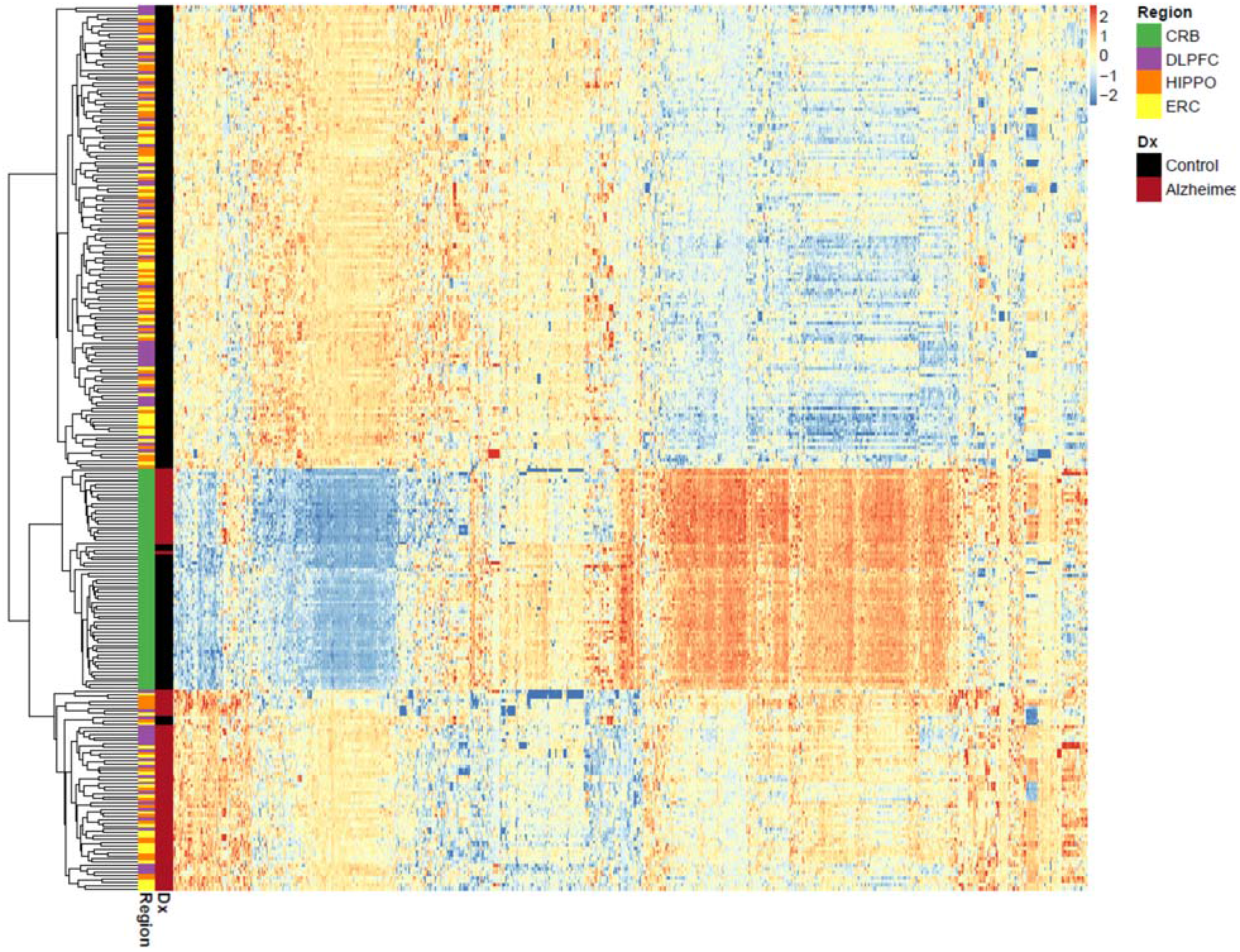
Heatmap of cross-region differentially methylated probes (DMPs). We adjusted beta values of 858 differentially methylated probes identified by the cross-region model by regressing out covariates (age, sex, ancestry and estimates technical variance), then Z-scaling the columns (DNAm sites). For hierarchical clustering and visualization purposes, Z-values less than the 1st percentile or greater than the 99^th^ percentile were set equal to these percentile Z-values (i.e. thresholded) to limit the effect of extreme values. Brain samples (rows) and DNAm sites (columns) were clustered using Euclidean distance (dendrogram not shown for columns).

We assessed the robustness of these findings using a series of sensitivity analyses. We found little effect of estimated neuronal composition on these results, as adjusting for the estimated proportion of neurons resulted in almost all DNAm sites (Sensitivity Model 1, 842/858, 98.1%) remaining FDR significant with highly correlated global t-statistics between both models (*r* = 0.954, p<2.2×10^−16^, **Figure S6A**). However, these results were somewhat expected given the lack of association between estimated neuronal composition estimates and diagnosis (p>0.068) in our dataset. Furthermore, differential DNAm does not appear to be driven by the greater burden of APOE risk in our AD samples: t-statistics were highly correlated with a model adjusting for APOE ε4 dosage (Sensitivity Model 2, *r* = 0.910, p<2.2×10^−16^, **Figure S6B**). Moreover, a slightly greater proportion of previously reported DMPs were consistent with the cross-region model than with the region-specific models of AD (**Table S5**). These analyses suggest cross-region DMP findings from our discovery model are not confounded by cell-type composition or APOE risk burden differences and previously reported DMPs are reasonably consistent.

### *Regional* DNAm differences between AD subjects and unaffected controls

Next, we tested for regional differences in DNAm that could relate to additional regulatory mechanisms [19] via differentially methylated regions (DMRs) using a bump hunting strategy that jointly tests neighboring DNAm sites for differential methylation [21]. We found four DMRs between AD cases and controls under a cross-region model (at family-wise error rate, FWER<0.05, **Table S6**). The most significant DMR was a hypermethylated region 1,136 base pairs (bp) long that overlapped an exon 57bp downstream from the *DUSP22* transcription start site (TSS) and spans 9 probes (FWER=0, **Figure S7A**). Post-hoc region-specific analyses revealed *DUSP22* hypermethylation in ERC (FWER=0.053), HIPPO (FWER=0.08), and DLPFC (FWER=0.088) but not in CRB (FWER=0.455). Another DMR overlaps the TSS of *ANKRD30B* (FWER=0.027, 511bp, 9 contiguous probes, **Figure 1E**). The third most significant region was less methylated and overlaps the promoter of *JRK* (384bp from TSS, FWER= 0.029, 5bp long, 2 contiguous probes, **Figure S7B**) and the fourth DMR overlaps *NAPRT* (FWER=0.047, 615bp long, 7 contiguous probes, **Figure S7C**). These results suggest a more limited role of regional changes in DNAm associated with AD.

**Figure 1E.**
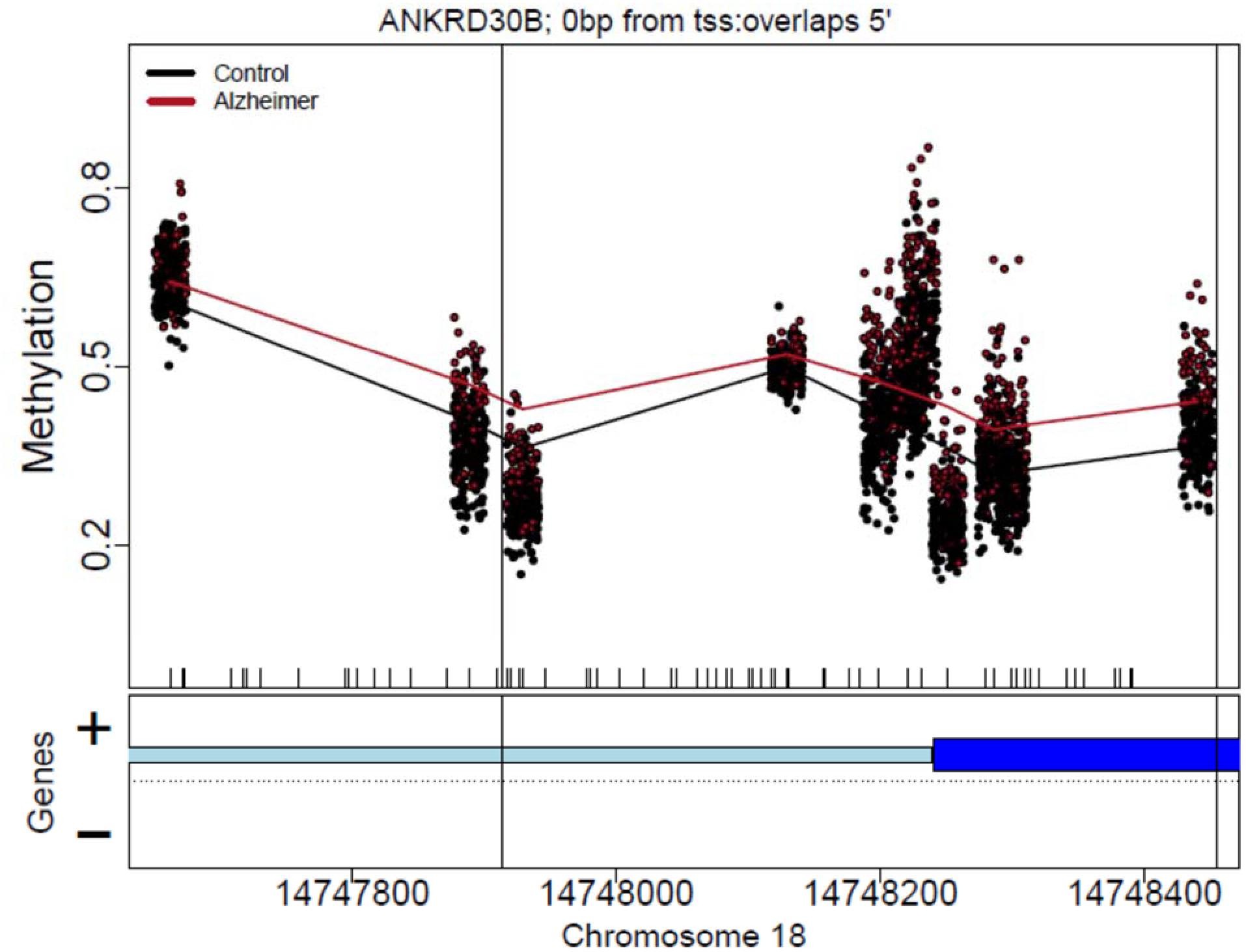
*ANKRD30B* differentially methylated region (DMR). A series of neighboring probes overlapping the transcription start site (TSS) of *ANKRD30B* are more methylated in Alzheimer’s disease (red) than in unaffected controls (black). *ANKRD30B:* Ankyrin Repeat Domain 30B

### Brain region-dependent differential methylation

Given that some brain regions, such as cerebellar cortex, are putatively less susceptible to AD pathology than other brain regions, we next assessed whether Alzheimer’s disease is associated with DNAm in a brain region-dependent manner (i.e. an interaction model, brain region by diagnosis). We found 11,518 DNAm sites with region-dependent AD effects (at FDR<5%, **Table S7**, **Figure S8**). The most significant region-dependent effect is seen for a DNAm site near *ANK1*, which is more methylated in AD subjects for DLPFC, ERC, and HIPPO but less methylated for CRB (cg11823178, p= 3.41×10^−21^, **Figure 2A**). Another example of a region-dependent effect can be seen for a CpG near *CSNK1G2* (cg01335597, p=3.04×10^−15^, **Figure 2B**). Region-dependent sites were mostly distinct from the cross-region DMPs identified above, though a subset of 130 sites was significant in both models (**Figure 2C**). These results demonstrate the power of combining data across multiple brain regions into joint statistical models and help further partition DNAm differences in AD by brain region.

**Figure 2A.**
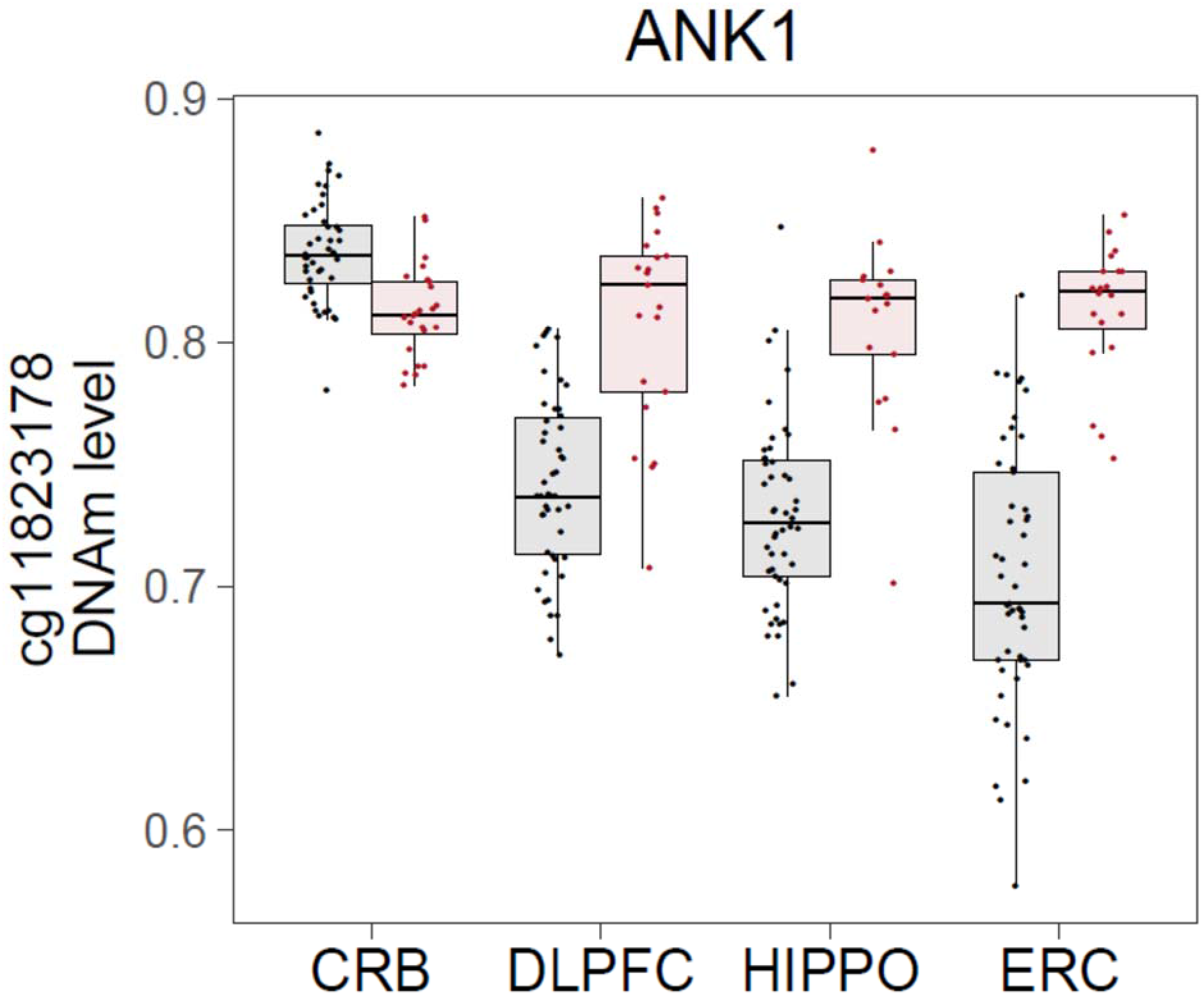
Example region-dependent DMP at cg11823178. Black points represent unaffected control samples and red points represent samples with a diagnosis of symptomatic Alzheimer’s disease (AD). The DNAm site cg11823178 is within 10kb of protein-coding gene *ANK1*. The effect of Alzheimer’s disease (AD) upon DNAm at this site is brain region dependent, with cortical brain regions, DLPFC, hippocampus (HIPPO), and entorhinal cortex (ERC), having a large difference between AD cases and unaffected controls, whereas little difference is observed in cerebellum (CRB). *ANK1:* Ankyrin 1

**Figure 2B.**
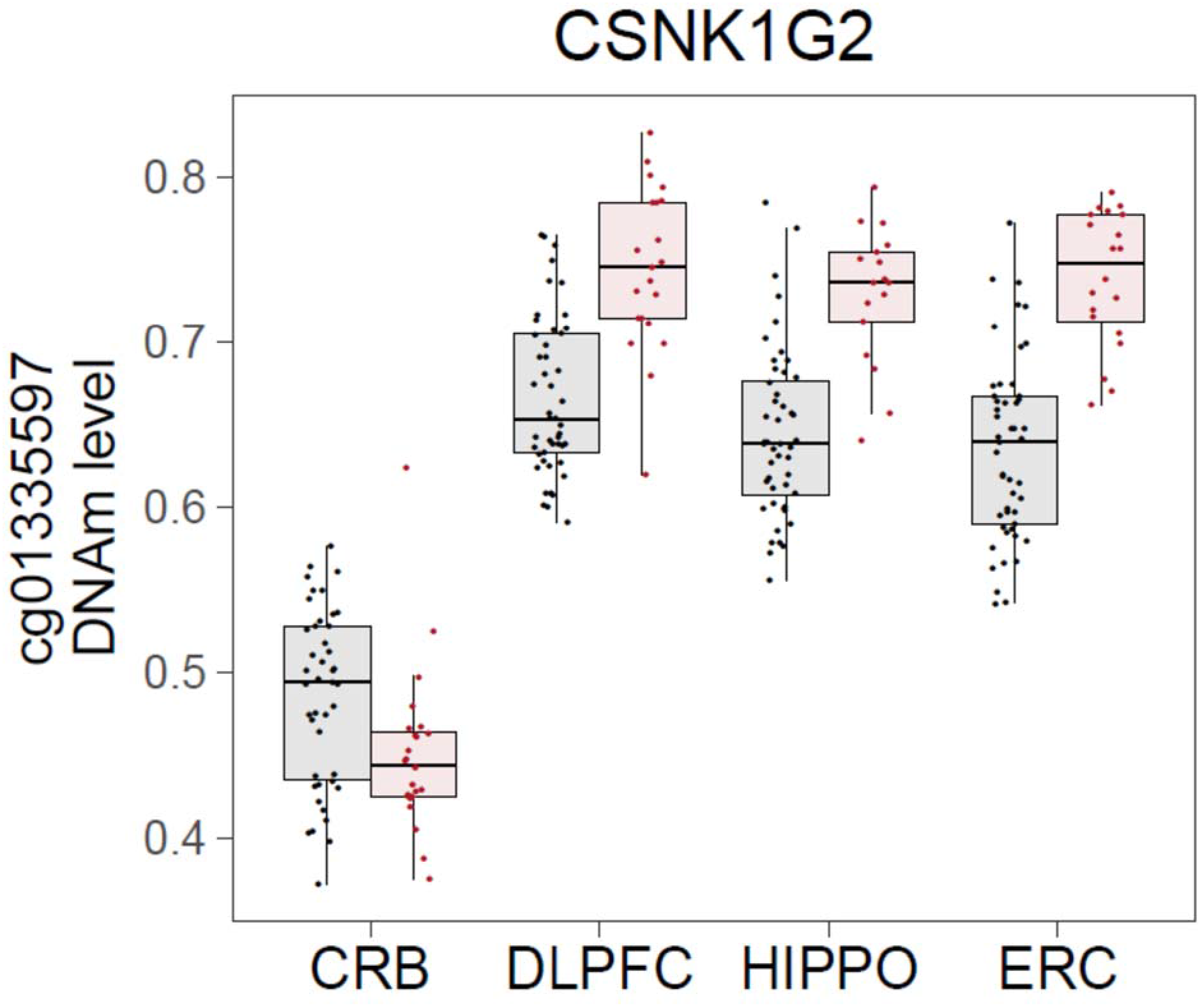
Example region-dependent DMP at cg01335597. The DNAm site cg01335597 is within 10kb of *CSNK1G2*, and is less methylated in CRB but more methylated in DLPFC, HIPPO, and ERC, in AD cases relative to unaffected controls. *CSNK1G2:* Casein Kinase 1 Gamma 2

**Figure 2C.**
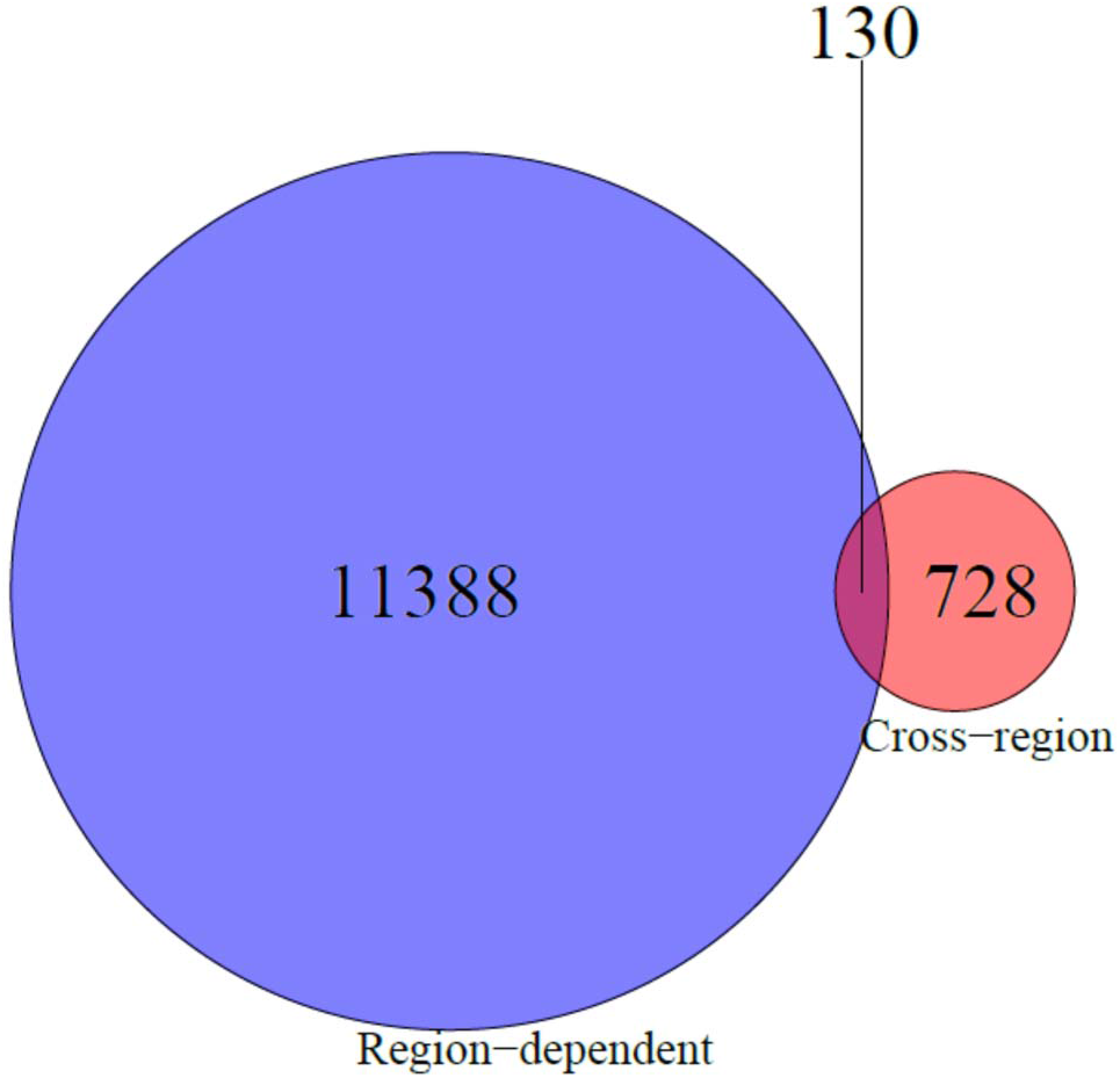
Venn diagram of cross-region and region-dependent sites. The number of overlapping and distinct cross-region and region-dependent DNAm sites is shown (at FDR<5%).

### Normal aging vs. AD-associated DNAm changes

Older age is a strong risk factor for Alzheimer’s disease [27] and indeed our AD donors were older than our controls (p=2.95×10^−8^). We, therefore, sought to disentangle the effects of normal aging on DNAm from the effects of AD on DNAm by assessing the association between DNAm and age in a sample of unaffected controls (N=187) and comparing these age-related associations to AD associations. Cross-region AD t-statistics were significantly associated with age-related t-statistics, but the absolute correlation was weak (Pearson’s *r*=0.089, **Figure S9A**), suggesting that on the genome-wide scale, cross-region DNAm differences in AD are generally distinct from the DNAm signature of normal aging. Nevertheless, significant AD-associated DNAm changes were highly enriched (Odds Ratio, OR=17.28, p<2.2×10^−16^) for similar age-related changes in unaffected controls (at p<1×10^−3^ and same directionality), further supporting the biological relevance of the most promising DNAm associations with AD with enrichment for similar changes during normal aging. In contrast, p-values from the region-dependent model were highly correlated with p-values from a region-dependent normal aging model (*r*=0.752, p<2.2×10^−16^, **Figure S9B**). We, therefore, focused further analyses on cross-region differential methylation effects, which appeared to be less susceptible to confounding by age (see **Results S4** for additional region-dependent analyses).

### Biological relevance of genes near differential DNAm

We next sought to better characterize the genomic and biological correlates of the DNAm changes identified under our cross-region model. Differentially methylated sites were enriched for CpG shelves and shores (**Figure S10**), suggesting these genomic features may be more dynamic in AD. Then we tested whether genes near differentially methylated sites were enriched in known pathways and ontologies while accounting for the biased distribution of CpGs per gene. Overall, DMPs were preferentially in close proximity to genes involved in processes related to cell-adhesion, immunity, and calcium binding (at FDR<5%, **Table S8A**). Further analysis suggests that hyper-methylated DNAm sites are driving these enrichments rather than hypo-methylated sites, which did not appear to be enriched in any particular process (**Tables S8B–C**). Thus, the gain of DNA methylation at these cross-region sites implicates genes involved in cell-adhesion, immunity, and calcium binding in Alzheimer’s disease.

### AD genetic risk loci are enriched for differential methylation

Given that AD risk has a strong genetic component, we investigated whether differential methylation was located preferentially within AD risk loci from a large GWAS meta-analysis [24]. While only a small number of differentially methylated sites were present in AD risk loci (5/858, 0.58%), it represents statistically significant enrichment when compared to the background overlap with risk loci (562/419,432, 0.13%; OR=4.37, p=0.00655, Fisher’s Exact Test, **Table S9**). This enrichment was fully driven by hypermethylated sites (N=5, OR=7.74, p=0.000588). When we relaxed our stringency for differential methylation (nominal p<0.01), the enrichment in GWAS loci remained statistically significant (OR=1.90, p=0.00423). These results indicate differentially methylated probes overlap putative AD genetic risk loci more than expected, which can hopefully be expanded upon and further refined through larger GWAS efforts in AD.

### Replication with an independent cohort

To assess the replicability of these cross-region DNAm differences, we re-analyzed processed, public AD DNAm data from four brain regions, DLPFC, ERC, CRB, and superior temporal gyrus (STG), under a similar model (N_control_=94 samples, N_AD_=239 samples from Lunnon et al. [31]; see Methods for further details). We found 28.0% of cross-region differentially methylated sites were consistent in this independent dataset (with p<0.05 and the same direction of effect; 240/858, OR=3.84, p<2.2×10^−16^, **Table S10A**). Notably, a site near *ANKRD30B* was more methylated in AD samples than controls (cg23703062, P_replication_=0.000355). These moderate replication rates are similar to those we observed previously (**Table S5**), suggesting reasonable concordance between AD DNAm differences from different brain cohorts.

### Using gene expression to functionally validate differential methylation

We next sought to more functionally validate differences in DNAm associated with AD, and tested for AD case-control gene expression differences. Here, we used RNA-seq data generated from a largely overlapping case-control series for the same multiple brain regions: HIPPO, DLPFC, ERC and CRB (N_donors_: 50 controls, 26 AD cases; N_samples_: 196 controls, 92 AD cases) to investigate expressed genes near a differentially methylated site (i.e. 645 DNAm sites corresponding to 772 genes within 10 kb; 218 DNAm sites were not within 10kb of a Gencode v25 annotated gene and were not considered for this analysis). The majority of genes near these sites were nominally differentially expressed between AD cases and controls in at least one brain region (p<0.05, 52.7% of genes, 407/772, **Table S11**). Interestingly *ANKRD30B*, which was implicated at both spatial resolutions (DMP and DMR), was under-expressed in AD patients compared to controls in ERC (log2 fold change LFC=−1.50, p=3.71×10^−5^) and HIPPO (LFC=− 1.70, p=0.00242) but not DLPFC (LFC=−0.434, p=0.198) or CRB (LFC=−0.304, p=0.665; **Figure 3A**). Other genes such as *WDR81* and *MYO1C* that were previously [8] implicated via differential DNAm were differentially expressed here (**Figures 3B–C**).

**Figure 3A.**
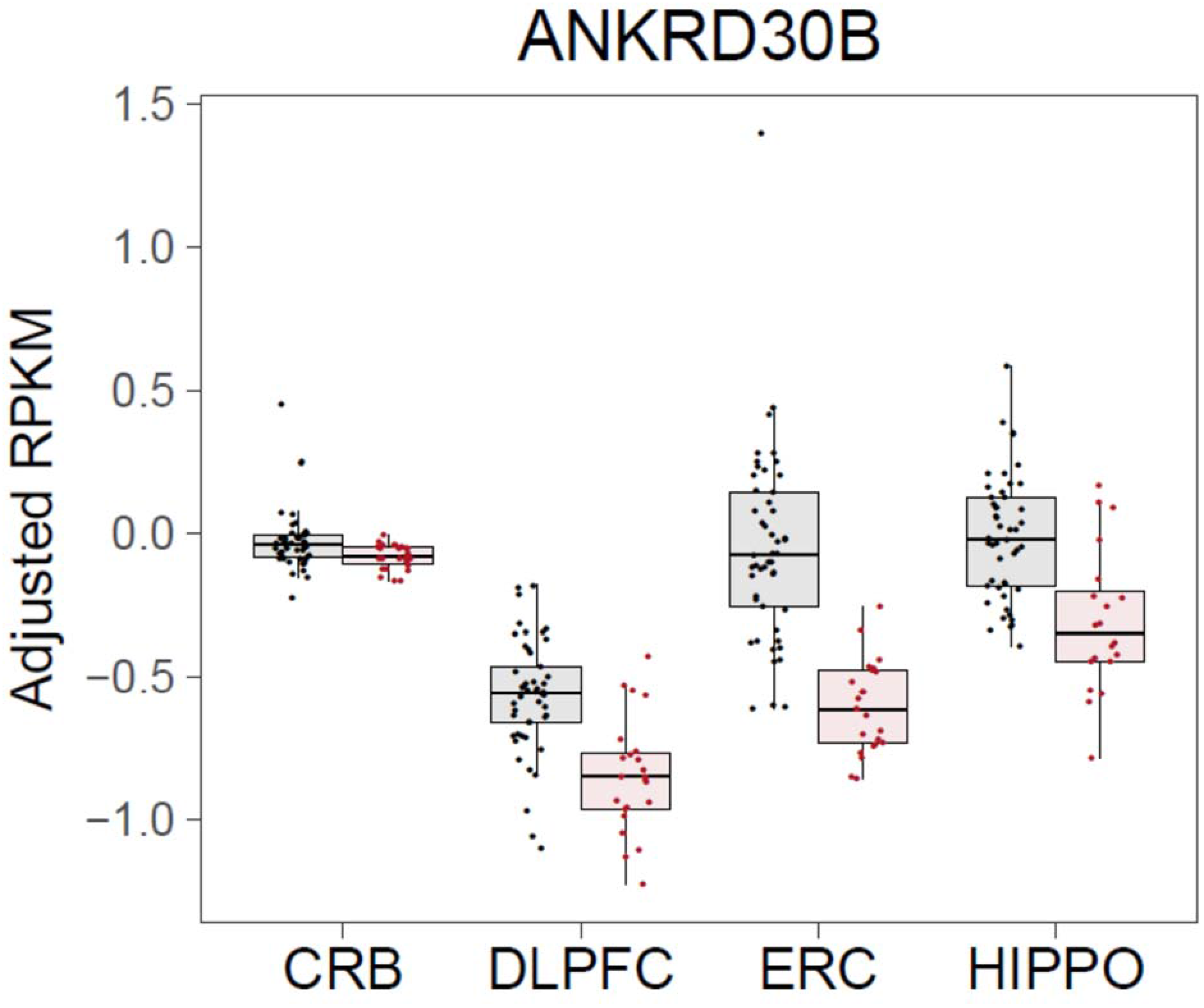
*ANKRD30B* differential expression. *ANKRD30B* is less expressed in AD samples compared to controls in cortical brain regions (DLPFC, ERC, and HIPPO), notably little difference between unaffected controls and AD cases is observed in CRB. *ANKRD30B:* Ankyrin Repeat Domain 30B

**Figure 3B.**
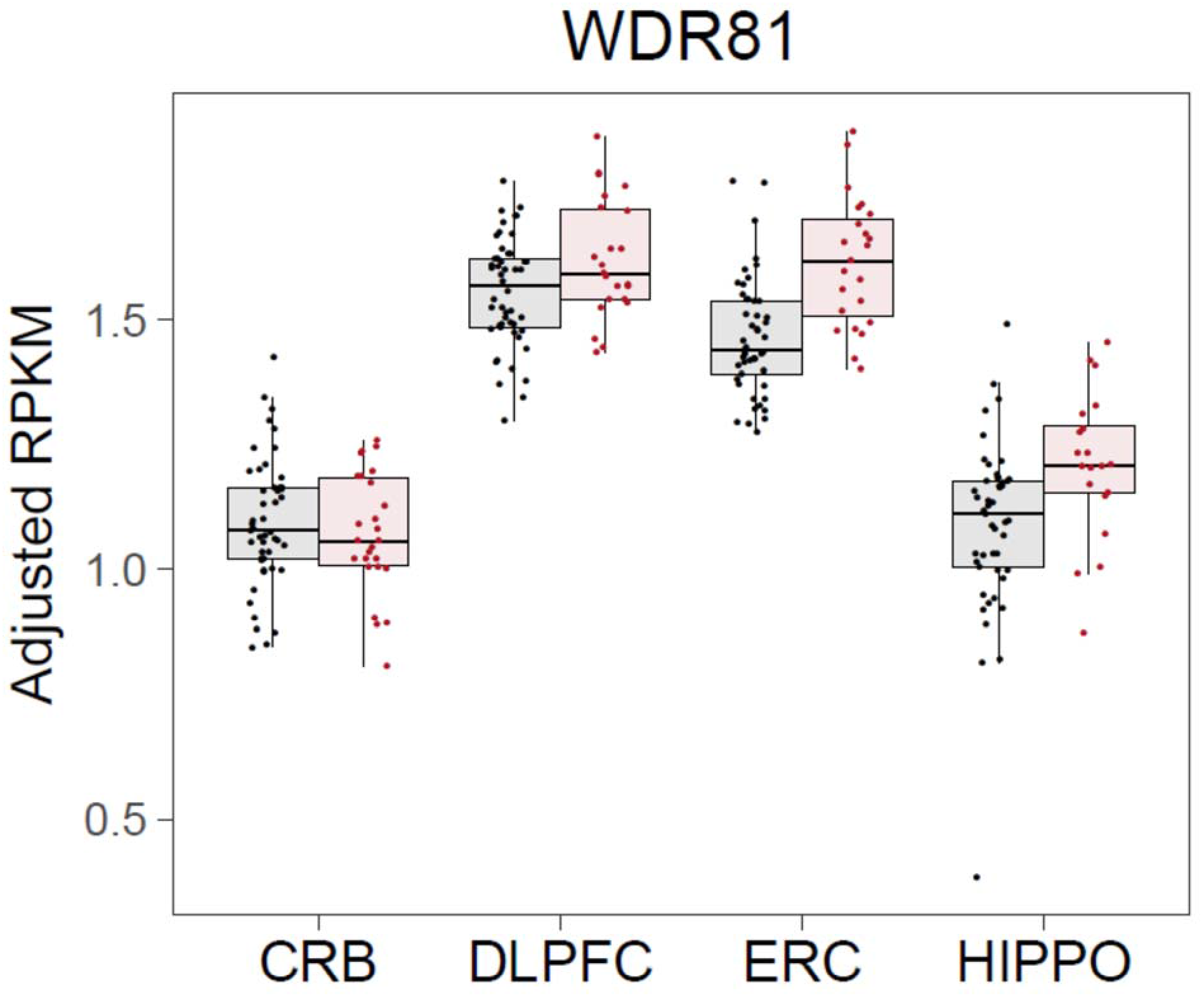
*WDR81* differential expression. *WDR81* is significantly more expressed in AD cases than in unaffected controls within ERC (CRB p= 0.915; DLPFC p= 0.111; ERC p= 0.00754; HIPPO p= 0.101). *WDR81:* WD Repeat Domain 81.

**Figure 3C.**
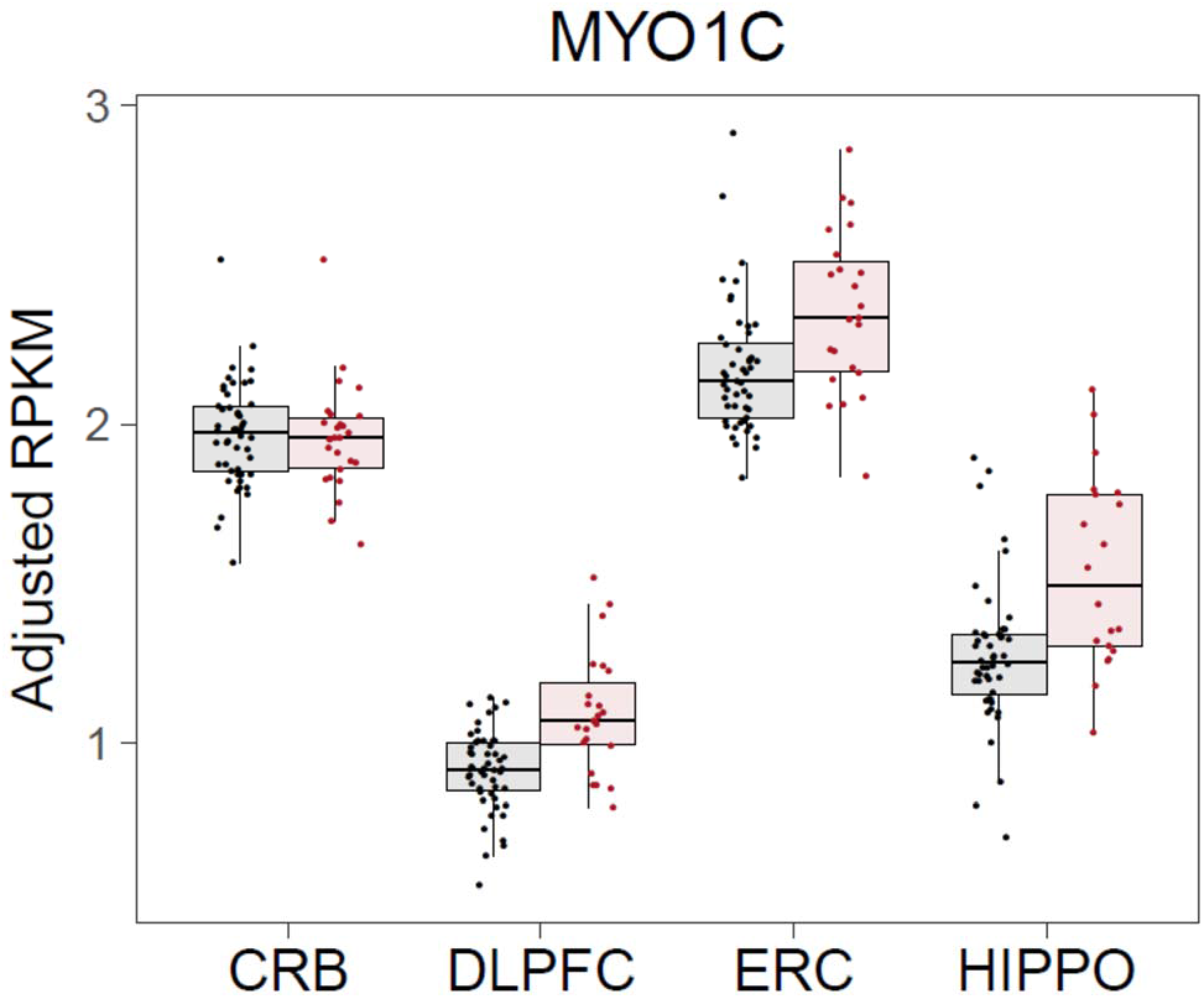
*MYO1C* differential expression. *MYO1C* is significantly more expressed in AD cases than in unaffected controls within cortical brain regions (CRB p= 0.826; DLPFC p= 0.00836; ERC p=0.0448; HIPPO p=0.0324). *MYO1C:* Myosin IC.

To further assess the functional link between DNAm and gene expression, we directly tested whether DNAm associates with gene expression for a subset of our samples that had both DNAm and RNA-seq data (N_donors_: 49 controls, 24 AD; N_samples_: 182 controls, 82 AD). We focused on the 407 genes with a nearby cross-region DMP and evidence of possible differential expression. We found DNAm associated with expression levels of 130 genes in at least one brain region (31.9%, at nominal p<0.05, **Table S12**). This integrated approach refined our list of candidate genes implicated by a nearby differentially methylated site (within 10 kb), improving the biological resolution of our epigenome-wide scan (*before:* 39.1% of CpGs with multiple genes, *after:* 8.23% of CpGs with multiple genes; *χ*^2^=55.069, non-parametric p-value = 1.6×10^−8^, 1×10^9^ bootstraps). Overall, DNA methylation was inversely correlated with gene expression (79/130, 60.8%, p=0.01406; using the strongest CpG-gene association), in line with the promoter-focused design of this microarray platform. Of note, hypermethylation of a site corresponding to *ANKRD30B* was associated with reduced gene expression in ERC, HIPPO, and DLPFC but not CRB (**Figure 3D**). While hypermethylation of a probe near *MYO1C* (cg14462670) associated with greater *MYO1C* expression in all four brain regions (**Figure S11A**), hypermethylation of a probe near *DUSP22* (cg11235426) associated with elevated gene expression only in DLPFC (**Figure S11B**). Together, these findings further implicate DNAm as a potential mechanism underlying dysregulated gene expression in Alzheimer’s disease.

**Figure 3D.**
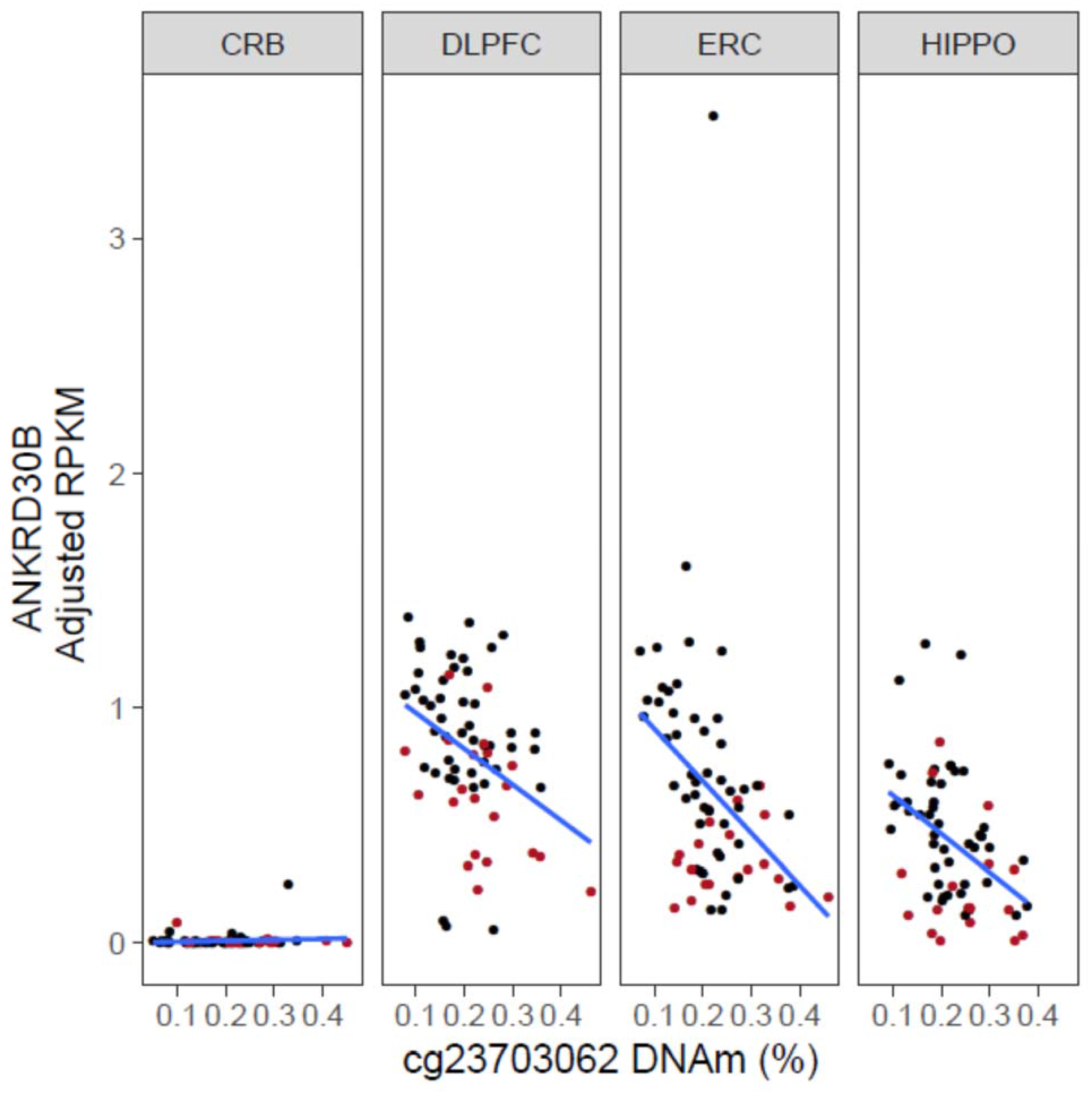
Greater DNAm at cg23703062 associates with reduced *ANKRD30B* expression in DLPFC, ERC, and HIPPO. Hypermethylation of DNAm site cg23703062, annotated to protein-coding gene *ANKRD30B*, associates with reduced gene expression in DLPFC, ERC, and HIPPO but not in CRB (CRB: p=0.910, β= 0.00755; DLPFC: p=0.0710, β= −0.719; ERC p= 0.0233, β= −1.25; HIPPO p=0.101, β= −0.792). *ANKRD30B:* Ankyrin Repeat Domain 30B

## Discussion

DNA methylation (DNAm) is an epigenetic factor that is disrupted in Alzheimer’s disease (AD), and the effects of this epigenetic disruption have already been linked to a few genes. To validate previously identified genes and potentially identify additional genes, we undertook a cross-brain region analysis of DNA methylation (DNAm) in Alzheimer’s disease (AD). Differential methylation implicated biological processes previously hypothesized to underlie AD pathology—cell-adhesion [30] and calcium ion homeostasis [32]—and were enriched in AD genetic risk loci [24]. By linking DNA methylation data with matching transcriptome data, we were able to prioritize 130 genes that were differentially expressed between cases and controls and for which DNAm associated with expression. Together these findings further validate several previous, prominent DNAm associations such as those with *ANK1* [8, 31] and *DUSP22* [43] (**Discussion S1**), and implicate novel epigenetically dysregulated genes.

Our study has several limitations. First and foremost, we cannot distinguish DNAm and gene expression differences that are causal in AD from those that relate to epiphenomena and secondary disease processes, such as neurodegeneration (**Discussion S2**). Indeed, the DNAm signature of other neurodegenerative diseases such as Lewy body dementia and Parkinson’s disease resembles that of Alzheimer’s disease [47]. Nevertheless, some of the DNAm differences reported here likely relate to AD etiology, because a subset of the differences was enriched within known late-onset AD genetic risk loci and correlated with gene expression changes. Further mechanistic studies to better untangle the timing of the co-occurring DNAm and expression changes will be necessary to more confidently determine which of these DNAm changes are causative versus an epiphenomenon in the brains of patients with Alzheimer’s disease.

In a similar vein, these DNAm changes can be regulated by other components of the genome and epigenome. For example, DNAm differences in AD can reflect underlying genetic risk, as was the case with a differentially methylated region overlapping the *PM20D1* promoter [46]. Likewise, DNAm changes may interact with other epigenetic factors that regulate gene expression [23]. Current human brain epigenome datasets now include post-translational histone modifications and higher order chromatin interactions, which have been shown to significantly alter gene expression both, within specific genetic loci and across the genome in human postmortem brain tissue [3, 40, 52]. Recent studies implicate widespread changes in the distribution of histone modifications in AD [11, 12]. While in some cases histone modifications may act independently of DNAm changes to generate AD risk [33], in other scenarios there is likely epigenetic cross-talk between DNA methylation and histone modifications [45]. Intriguingly, whereas a study of H4K16 acetylation identified AD-related differences weakly inversely associated with age-related changes [39], we observed the opposite correlation with DNAm, albeit the association was weak. The canonical roles of DNA methylation as a transcriptionally repressive epigenetic mark and histone acetylation as an activating one may reconcile these conflicting patterns. Disruption of multiple epigenetic factors may converge on transcriptional networks [36, 38, 54] that are disrupted in Alzheimer’s disease.

Our study suggests disrupted DNAm plays a role within this complex gene regulatory framework. We identified 130 genes that had evidence of differential expression between cases and controls and for which expression associated with nearby DNAm that differed in AD. These differentially expressed genes may involve cell type-specific dysregulation. For instance, a recent study of cell-sorted cortical tissue found DNAm differences existed in both neurons and glia that associated with AD neuropathology (Braak stage) [10]. Likewise, a study of laser captured brain tissue suggests microglia are responsible for differential expression of the epigenetically dysregulated gene, *ANK1* [35]. Understanding the transcriptional output of these epigenetic changes in AD across multiple brain regions, which have different susceptibilities to AD neuropathology, will be a challenging, but rewarding endeavor. Our publicly available dataset contributes unique matched RNA-seq and DNAm data that can be used for development of genomic integration statistical methods.

Studying multiple brain regions is crucial to convert knowledge of epigenetic changes into insight of molecular risk mechanisms, because each region has a distinct cellular composition and regulatory landscape that contributes to neurophysiological function [7, 14, 20, 37, 44]. By studying the epigenome and transcriptome of four brain regions in parallel, three cortical regions susceptible to AD, and cerebellum, a region that is relatively protected, we were able to cast a wide net to capture AD-related differences. While we focused upon “cross-region” differential methylation—changes concordant across multiple brain regions—we also identified and characterized a large number of region-dependent differentially methylated sites. Moreover, even if differential methylation is shared across cerebellar and cortical brain regions, the gene expression correlates may not be. For example, *ANKRD30B* is hypermethylated in all four brain regions but is not expressed at observable levels in cerebellum. By studying the unique epigenetic profiles and their transcriptional correlates of multiple brain regions, we may glean deeper insight into the molecular mechanisms underlying differential susceptibility of brain regions to AD neuropathology.

In summary, we used paired DNA methylation and transcriptome data from four brain regions to link DNAm differences in AD to local gene dysregulation. These results support the role of DNAm as an epigenetic mechanism underlying gene dysregulation and support novel genes involved in AD. More generally, they illustrate the value of integrating epigenetic and transcriptomic data to study complex disease.

## Methods

### Postmortem brain tissue dissections

DLPFC (Brodmann areas 9 and 46), hippocampal formation, entorhinal cortex at the level of the anterior hippocampus and cerebellar cortex were dissected from frozen postmortem brains using a hand-held visually guided dental drill (Cat #UP500-UG33, Brasseler, Savannah, GA) as previously reported [28]. In addition to demographic matching between Alzheimer’s disease control subjects, the subjects included in the control group had no clinical or neurological history, or history of alcohol or substance abuse, or positive toxicology screens for illicit substances.

### Alzheimer’s neuropathology diagnosis

All postmortem brains of subjects (controls and Alzheimer’s disease) have been sampled for neuropathology and specific Alzheimer’s lesions: beta-amyloid plaques, neurofibrillary tangles and tau-positive neurites. Samples were also genotyped for the apolipoprotein E gene *(APOE)*. The sampled tissue sections were fixed in 10% buffered formalin and paraffin embedded into blocks for microscopic analysis (10 micron thickness). Sections included the superior frontal gyrus, middle and superior temporal gyri, inferior parietal cortex, occipital cortex, amygdala, hippocampus and entorhinal cortex, anterior thalamus, midbrain, pons, medulla, and cerebellum (including cerebellar cortex and deep cerebellar nuclei). Sections were silver-stained using the Hirano method (Yamamoto and Hirano, 1986) and immune-stained using antibodies against ubiquitin, phosphorylated anti-tau (PHF-1) and beta-amyloid protein (6E10). Microscopic preparations were examined using conventional light microscopy. Alzheimer’s neuropathology ratings include the Braak (Braak H, Braak E, 1991) staging schema evaluating tau neurofibrillary tangle burden, and the CERAD scoring system (Mira SS et al., 1991) as a measure of senile plaque burden (neuritic and diffuse). An Alzheimer’s likelihood diagnosis was then performed based on the published consensus recommendations for postmortem diagnosis of Alzheimer’s disease (Hyman BT, Trojanowski JQ, 1997) as in prior publications (Conejero-Goldberg C et al., 2015).

### Processing DNA methylation data

Human Methylation 450k (HM450k) arrays were run as specified by the manufacturer upon DNA extracted from each brain region. After generating array data for 398 postmortem brain samples, resulting idat files were imported, then rigorously preprocessed with minfi [1]. We did not observe batch effects with several different quality control metrics. After removing low-quality samples (N=5), we normalized the data with stratified quantile normalization [1]. We removed 64,660 probes that met one or more of the following exclusion criteria: (a) poor quality (b) cross-reactive (c) included common genetic variants (d) mapped to a sex chromosome (e) did not map to hg38. We dropped 13 samples one or more of the following reasons: (i) DNAm predicted sex did not match phenotypic sex (ii) 450k genotype clustered inappropriately (iii) 450k genotype did not match SNP-chip genotype (iv) clustered inappropriately on principal component analysis. After these conservative quality control steps, 420,852 high-quality probes available for 377 samples remained for further analysis. We estimated cell-type compositions using deconvolution algorithm [17] with a flow-sorted DLPFC reference [13] and estimated age acceleration with Horvath’s clock[16]. For additional details about DNAm processing see Supplemental Methods (**Figures S12-S17**).

### Brain-region stratified DNAm analyses

We tested for differences between AD cases and controls in predicted neuronal proportion and age acceleration, stratified by brain region, with linear models. To test the association between DNAm and AD across the epigenome, we stratified samples by brain region, then tested beta values from 420,852 probes for differential methylation between Alzheimer’s disease cases and controls using limma [42]. We included age, sex, ancestry, and negative control principal components 1 and 2 as covariates in our model and controlled for multiple testing by using the false-discovery rate (FDR).

### Cross-region differential methylation analysis

In our “cross-region” model for AD case-control differences (N=269 samples), we tested beta values from 420,852 probes for differential methylation between Alzheimer’s disease cases and controls using limma[42] with the model:

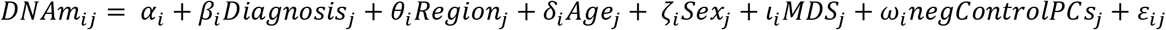

for DNAm site *i* and donor *j* and binary disease status *Diagnosis_j_* adjusting for brain region, age, sex, the first multidimensional scaling (MDS) component of genotype data (ancestry), and the first two negative control principal components (PCs). Afterward, we tested an interaction model, i.e. an F-test of the estimated AD effect from each brain region, while adjusting for the same covariates as above. Both of these models incorporated multiple brain regions from each donor, and thus repeated measures, so we used the generalized least squares technique to estimate statistical parameters to appropriately account for the repeated biological measures. We estimated a consensus correlation with limma’s *duplicateCorrelation* function in which donors are blocks and different brain regions are repeated observations. Then, we incorporated this consensus correlation while estimating statistics for each DNAm site using the generalized least squares technique via limma’s *lmFit* function. We performed four sensitivity analyses to assess how robust our results were to potential confounds (see Supplemental Methods for additional details).

### Differential methylated region analysis

To assess whether regions of the genome were differentially methylated between Alzheimer’s and controls, we used bumphunter with the same models as above [21]. We implemented the bumphunter method with the minfi Bioconductor package’s *bumphunter* function using 1000 bootstrap iterations, a cutoff of 0.05 for defining continuous probes, smoothing done with the *locfitByCluster* function, and otherwise default parameters [1]. We controlled for multiple testing by using the family-wise error rate (FWER) to determine statistical significance.

### Differential expression of DMP genes and association with DNAm

DNA methylation sites were annotated to all Gencode v25 genes within 10kb upstream or downstream (20kb region) using the GenomicRanges Bioconductor package [25]. These genes were then tested for corresponding AD-case control differential gene expression. Analyses were stratified by brain region and used 196 control and 92 AD RNA-seq samples processed using a standardized pipeline. We normalized counts for 25,587 Gencode annotated genes that were expressed in at least one brain region with limma’s *voom* function then tested for AD case-control differences under an empirical Bayesian framework [42] while adjusting for RNA Integrity Number (RIN), age, sex, race, mitochondrial mapping rate, and gene assignment rate.

We tested then the association between DNAm at each differentially methylated site and log-transformed, normalized gene expression (log2[RPKM+1]) for all Gencode (v25) annotated genes within 10kb with at least partial evidence of differential expression (nominal p<0.05 in at least one brain region). This analysis used 185 control and 85 AD samples with matching DNA methylation and RNA-seq data. We implemented this association test between DNAm and gene expression in R using a linear model (*lm* function) that adjusted for the covariates above and was stratified by brain region.

## Ethics approval and consent to participate

Post-mortem human brain tissue was obtained by autopsy primarily from the Offices of the Chief Medical Examiner of the District of Columbia, and of the Commonwealth of Virginia, Northern District, all with informed consent from the legal next of kin (protocol 90-M-0142 approved by the NIMH/NIH Institutional Review Board).

## Consent for publication

Not applicable.

## Availability of Data and Materials

Raw and processed data will be made publicly available in accordance with editorial policy. The computer code used to generate results is available on GitHub: https://github.com/LieberInstitute/Alzheimers DNAm

## Author contributions

S.A.S. – performed analyses and led the writing of the manuscript

R.A.B. – assessed neuropathology, contributed to biological study design and interpretation, and writing of the manuscript

L.C.T. – processed RNA-seq data and contributed to its analysis

R.T. – performed APOE4 genotyping

J.H.S. – performed data generation

A.D. – performed clinical reviews

J.W. – contributed to the analysis of the DNAm data

D.R.W. – contributed to the study design, interpretation of the results, and writing of the manuscript

T.M.H. – performed tissue dissections, contributed to the study design, interpretation of the results, and writing of the manuscript

J.E.K. – oversaw tissue collection and contributed to the study design, interpretation of the results, and writing of the manuscript

A.E.J, V.S.M. – co-led the study, including the design, statistical analyses, interpretation, and writing of the manuscript

## Supplemental Appendix

### Supplemental Results

#### S1. Cell-type composition

Although AD is associated with neuronal loss across multiple brain regions[1–3], there is mixed evidence for different proportions of neurons in AD brain when using cell-type specific signatures of DNA methylation (DNAm)[4–6]. We used a previously published, deconvolution algorithm[7] to estimate the proportion of neuronal cells in each of our samples using DNAm cell-type signatures (see Methods). The estimated proportions of neurons was not significantly different between AD and controls in any brain region, though we observed a trend towards reduced neuronal composition in DLPFC, hippocampus, and entorhinal cortex (p>0.05, linear regression, **Figure S2**).

#### S2. Epigenetic age acceleration

Epigenetic age acceleration, a putative biomarker of more rapid biological aging than expected, was previously reported to be weakly correlated with AD pathology in the DLPFC[8]. We estimated the epigenetic age of our samples using a validated method (“Horvath’s clock”)[9]. Epigenetic age was highly correlated with chronological age across all regions confirming that the estimation was successful (mean *r*=0.812, p<2.2×10^−16^, **Figure S3A**). Epigenetic age acceleration, however, was not significantly associated with AD diagnosis in DLPFC or in any other brain region, though we observed a trend towards acceleration in DLPFC for AD subjects (p>0.05, t-test, **Figure S3B**).

#### S3. Region-stratified differential methylation analysis

To assess DNAm differences associated with AD in a brain region-specific manner, we undertook an AD case-control analysis stratified by brain region. We tested 420,852 probes for differential methylation between unaffected controls (N≤49) and donors with Alzheimer’s disease (AD, N≤24). With this region-specific, case-control analysis, we identified four differentially methylated probes (DMPs, at false discovery rate, FDR<5%): three hypermethylated DMPs in dorsolateral prefrontal cortex (DLPFC) and one in entorhinal cortex (ERC) (**Figure S4, Table S2**). No genome-wide significant DMPs were found in hippocampus (HIPPO) or cerebellum (CRB).

Two of the DLPFC DMPs tagged genes that were previously reported[10] as hypermethylated: *MYO1C* (cg05417607, p=1.19×10^−7^, change in DNAm, Δ=0.065, **Figure S4A**) and *SPG7* (cg03169557, p=1.74×10^−7^, Δ=0.082, **Figure S4B**). The third DMP corresponds to a putatively novel association implicating *MARCKS* (cg10474881, p=1.35×10^−7^, Δ=0.046, **Figure 4C**). The ERC DMP replicated a previously identified association with the gene *ANK1*[4, 5] (cg05066959, p=1.04×10^−7^, Δ=0.13, **Figure 4D**). Notably, the three previously reported DMPs (*MYO1C, SPG7, ANK1*) were highly significant (p<1×10^−5^) in at least one other brain region with directional consistency, whereas the association with *MARCKS* lacked similarly convergent evidence (**Table S2**).

Furthermore, reported associations from De Jager et al.[4] and Lunnon et al.[5] were moderately replicable in our data when comparing the same region (10%–40%, **Table S3**; at p<0.05 and same directionality of effect, see Methods for further details). Interestingly, we observed comparable replication for De Jager’s DLPFC results even when we used different regions (HIPPO=40.6%, ERC=36%). These findings suggest that DNAm changes associated with Alzheimer’s disease are most replicable when using the same tissue, though some of these DNAm differences may also be shared across other brain regions.

#### S4. Region-dependent AD differential methylation

We further explored the biological correlates for the set of 11,518 DNAm sites with brain region-dependent DNAm differences (i.e. brain region by diagnosis interaction). For this set of sites, post-hoc statistics show that the effect of AD in the cerebellum is negatively correlated with the effect in AD within the other three brain regions (**Figure S9**), suggesting the DNAm differences in Alzheimer’s disease in the CRB differs strongly those within ERC, HIPPO, or DLPFC. Genes near (within 10kb) region-dependent differentially methylated sites were also enriched for pathways related to cell adhesion (GO_BP_, “regulation of cell adhesion”, p=7.30×10^−5^) and calcium signaling (KEGG, “calcium signaling pathway”, p=0.000401), although interaction CpGs were most highly enriched for biological processes related to development, e.g. “anatomical structure morphogenesis” (p=2.86×10^−15^) and “developmental process” (p=1.19×10^−14^, **Table S8D**). Region-dependent sites were not preferentially located within AD risk loci, at either false-discovery rate significance (p=0.897, Odds Ratio, OR=1.03) or nominal significance (at p<0.01; OR=1.18, p=0.325, **Table S9**). We found 25.2% of our brain region-dependent CpGs were consistent in an independent dataset (nominal p<0.025, 2,908/11,518, OR=5.78, Fisher’s p<2.2×10^−16^, **Table S10B**; data from Lunnon et al[5]). These results suggest region-dependent differentially methylated sites are a functionally distinct from the cross-region sites.

### Supplemental Discussion

#### S1. Differentially methylated regions

Our most significant differentially methylated region (DMR) overlapped *DUSP22* and this association was significant (FWER<10%) in dorsolateral prefrontal cortex (DLPFC), hippocampus (HIPPO), and entorhinal cortex (ERC). Interestingly, a previous study[10] found hyper-methylation of the *DUSP22* promoter in hippocampus and showed DUSP22 protein affects tau phosphorylation and CREB signaling. A DMR overlapping *DUSP22* has also been reported in the superior temporal gyrus of AD brains but with the opposite direction of effect (hypomethylation in AD)[6]. These results suggest *DUSP22* is hypermethylated in multiple cortical brain regions beyond the hippocampus, such as entorhinal cortex and dorsolateral prefrontal cortex.

Three other DMRs were identified which have not been previously reported in the context of AD, to our knowledge. One of these novel DMRs overlaps the transcription start site (TSS) of Ankyrin repeat domain 30B (*ANKRD30B*), a protein-coding gene of unknown function which has been linked to breast cancer[11] and rare neurodevelopmental disorders—Williams syndrome and 7q11.23 duplication syndrome[12]. Another DMR overlaps the promoter of *JRK* and is 384 bp upstream of the TSS; genetic studies of this gene provide mixed evidence for its role in childhood epilepsy {7550318; 11463517; 10510981}. The fourth DMR is 11 bp downstream from the transcription start site of *NAPRT*, a gene which plays role in bioenergetics, particularly in synthesizing NAD+, the abundance of which is altered in late-onset AD{29070876}. Thus, disruption of the genes near the four differentially methylated regions have been previously linked to brain dysfunction, though they do not appear to fall into a single coherent pathway.

#### S2. Cell-type composition

We also cannot fully disentangle changes in cell-type composition from the differential methylation signals. Although we did not find differences in the estimated proportion of neurons with the *in silico* deconvolution algorithm and a sensitivity analysis accounting for these estimates did not change our conclusions as others have seen[4],[5], loss of neurons{**7286917; 7916070; 7058341; 8699259**} and increased neuroinflammation{**25792098; 22315714; 10858586**} are hallmarks of Alzheimer’s disease neuropathology. This discrepancy may arise due to limitations of the *in silico* deconvolution or from postmortem brain sampling which may include both healthy and diseased tissue. Here, cell-type specific approaches to DNAm profiling will be crucial for resolving tissue composition heterogeneity and for unmasking cell-type specific neuropathological differences.

### Supplemental Methods

#### Preprocessing and normalization

idat files from 398 samples were imported with minfi[13] for rigorous preprocessing. Principle components (PCs) were computed upon the log-transformed (log_2_[x+1]) red and green negative control probes (N=1,226). The first two negative control PCs together captured ~49.1% of the variance and were used later as covariates to control for technical variation such as plate effects, which is considered a “batch effect” (**Figure S13**). Samples with outlying distributions of methylation/unmethylation intensity were removed (N=5, **Figure S14**). Notably, we did not observe systemic variability of the data quality associated with disease or brain regions, suggesting “batch effects” were unlikely to confound these biological effects of interest. We then normalized the data with stratified quantile normalization[13] which improved the distribution of beta values (**Figure S15**).

#### Filtering probes

We removed probes which were of low quality, potentially affected by common SNPs, cross-reactive, mapped to sex chromosomes, or mapped ambiguously to hg38. 2,186 probes were dropped because they had a detection P-value greater than 0.05 in over 5% of our samples. 16,864 probes were dropped because they contained a common single nucleotide polymorphism (minor allele frequency, MAF>1%) at either single base extension sites or the target CpG. We dropped 27,386 probes that Chen et al. reported as cross-reactive[14], and removed 10,258 probes that mapped to sex chromosomes (chrX and chrY). Lastly, we excluded 7,966 probes which we were unable to confidently map to hg38. After removing these 64,660 probes, 420,852 high-quality probes available for 393 samples remained for further analysis.

#### Dropping samples

We identified and removed samples where methylation predicted sex did not match phenotypic sex (N=4, **Figure S16**), and samples whose genotype determined by 65 SNPs present on HM450k array clustered incorrectly (N=2, **Figure S17**). Samples whose methylation genotype did not match their SNP-chip genotype were also dropped (N=7). We removed three samples that were labeled as cerebellum but clustered with cortical samples in the first principal component of DNAm probe beta values (**Figure S18**). After removing these 16 samples, 377 high-confidence samples remained for further analysis.

#### Cell type composition

To estimate the proportion of NeuN+ cells, we used a modified version of the algorithm described in Houseman et al. 2012[7] implemented in minfi with a flow-sorted DLPFC reference profile methylation {**Jaffe AE and Kaminsky ZA (2017). FlowSorted.DLPFC.450k: Illumina HumanMethylation data on sorted frontal cortex cell populations. R package version 1.14.0; 23426267**}. After predicting the cell-type composition, we tested for AD case-control differences in neuronal proportions using a linear model separately for each brain region while adjusting for age, sex, and MDS 1 (a measure of ancestry).

#### Age acceleration

Epigenetic age was predicted with Horvath’s clock as recommended[9]. We defined epigenetic age acceleration as the residuals from the linear model between chronological age and DNAm age. Epigenetic age acceleration was calculated separately across all four brain regions, and we then tested for different epigenetic age acceleration between controls and AD samples with a two-tailed t-test.

#### Region stratified epigenome-wide scan

We stratified samples by brain region, then tested beta values from 420,852 probes for differential methylation between Alzheimer’s disease cases and controls using limma[15] with the model:

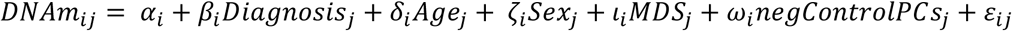

for CpG *i* and donor *j* and binary disease status *Diagnosis_j_*, adjusting for age, sex, the first MDS component (ancestry), and negative control principle components 1 and 2 for each donor. Multiple testing was controlled for in this analysis and those following (unless otherwise specified) by using false-discovery rate (FDR) adjusted p-values to determine significance.

#### Sensitivity analyses

We performed three sensitivity analyses to assess how robust our results were to the potential confounds. In the first analysis (Sensitivity Model 1), we included the predicted proportion of NeuN+ (neuronal) cells as a covariate. In the second analysis (Sensitivity Model 2), we included APOE ε4 allele dosage as a covariate in our model for AD case-control differences. In another secondary analysis, we estimated age-related effects upon DNAm using only unaffected controls (N=190) for 420,852 probes, while adjusting for sex, ancestry, brain region, and negative control PCs 1 and 2 (accounting for repeated measures as described in the main text). Then, we compared these effects sizes to those from the AD case-control model.

#### Replication with independent results

To assess the replicability of previous studies, we downloaded published results from two landmark epigenome-wide association studies of AD[4, 5] and checked whether their DMP findings replicated in our data. Here, we defined “replicable” as p<0.05 and the same direction of effect. Hence, “percent replicated” is defined as:

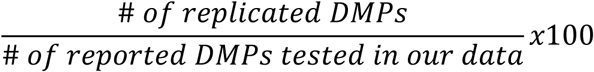

Conversely, we assessed the replication of significant DMPs discovered through cross-region and region-dependent analyses. We analyzed processed data from three cortical regions and cerebellum from Lunnon et al.^10^ (N_control_=94 samples, N_AD_=239 samples, GSE59685) with a generalized least squares model using the diagnosis provided and including brain region as a covariate.

#### Enrichment tests

Probes were mapped to Entrez IDs and we tested for gene set enrichment of DMPs in GO and KEGG ontologies using an adaptation of the *gometh* function from the missMethyl package[16]. The *gometh* function reduces potential spurious enrichment signals arising from the non-uniform distribution of CpGs per gene by accounting for the number of probes mapping to each gene. The 420,852 probes we tested for differential methylation were used as the background. When evaluating changes in the distribution of CpG genomic density features (i.e. island, shore, shelf, open sea), we used a chi-square test of the 2×4 contingency table. Further post-hoc tests were performed with Fisher’s exact test (2×2 contingency tables with other non-tested categories grouped together). These analyses were done using the set of FDR significant probes by AD case-control status.

To test whether differentially methylated probes were enriched in AD genetic risk loci, we first downloaded the 21 genetic risk markers available from the largest meta-analysis published at the time of analysis (Table 2 of Lambert et al.)[17]. We then constructed linkage-disequilibrium (LD) blocks by identifying proxy markers in LD (R^2^>0.6) with the 21 index marker using rAggr (1000Genomes Phase3 hg19 reference, African + European populations, MAF>0.001, distance<500kb, HW p-value cutoff=0, at least 75% genotyped; http://raggr.usc.edu/). Cumulatively, these LD blocks covered 2429.18kb of the genome. We overlapped the coordinates of these LD blocks with those of our DNAm probes using the GenomicRanges Bioconductor package[18] and identified 567 tested probes that were within the LD boundaries for 19 of these AD risk loci. Enrichment was calculated using Fisher’s Exact Test of the 2×2 contingency table of probes differentially methylated vs. within AD risk loci.

### List of Supplemental Figures

**Figure S1**. Distribution of neuropathology, cognitive performance, age, and brain mass in our dataset.

**Figure S2**. Estimated proportion of neurons in AD and control samples across the four brain regions studied.

**Figure S3A**. Epigenetic age is highly correlated with chronological age across the four brain regions studied.

**Figure S3B**. Distribution of epigenetic age acceleration in AD and controls for the four brain regions studied.

**Figure S4**. Four significant differentially methylated probes (DMPs) were identified under region stratified models.

**Figure S5**. Post-hoc effect sizes from region-specific models for cross-region DMPs are highly correlated across all brain regions studied.

**Figure S6A**. Adjusting for cell-type composition increases t-statistics of AD case-control differences.

**Figure S6B**. T-statistics from model adjusting for APOE4 burden are highly correlated with the original model.

**Figure S7A**. Differentially methylated region overlapping *DUSP22*.

**Figure S7B**. Differentially methylated region nearest to *JRK*.

**Figure S7C**. Differentially methylated region nearest to *NAPRT*.

**Figure S8**. Post-hoc scatter plots of effect sizes from four brain regions for interaction CpGs.

**Figure S9A**. T-statistics from cross-region AD model versus t-statistics from cross-region aging model.

**Figure S9B**. T-statistics from region-dependent AD model versus t-statistics from region-dependent aging model.

**Figure S10**. Hypermethylated CpGs enriched for island shelves.

**Figure S11A**. DNAm associates with *MYO1C* expression across all four brain regions.

**Figure S11B**. DNAm associates with *DUSP22* expression only in DLPFC.

**Figure S12**. Negative control PCs 1 and 2 associate with batch (sample plate).

**Figure S13**. QC plot.

**Figure S14**. Distribution of beta values before and after stratified quantile normalization (SQN). **Figure S15**. Four potential sex swaps were removed.

**Figure S16**. Two samples clustered incorrectly based on genotype. These samples were removed prior to further analysis.

**Figure S17**. Principal component analysis of processed data.

### List of Supplemental Tables

**Table S1**. Characteristics of samples studied.

**Table S2**. Four significant differentially methylated probes (DMPs) under region-specific models.

**Table S3**. Region-specific model concordance with previously reported findings from De Jager et al.[4] and Lunnon et al.[5]

**Table S4**. 858 differentially methylated probes (DMPs) under a cross-region model.

**Table S5**. Cross-region model concordance with previously reported findings from De Jager et al.[4] and Lunnon et al.[5]

**Table S6**. Results for three significant cross-region differentially methylated regions (DMRs) and corresponding post-hoc region-specific statistics.

**Table S7**. 11,518 differentially methylated probes (DMPs) under a region-dependent model.

**Table S8A**. Gene ontology results for 858 cross-region significant probes.

**Table S8B**. Gene ontology results for cross-region significantly hypomethylated probes.

**Table S8C**. Gene ontology results for cross-region significantly hypermethylated probes.

**Table S8D**. Gene ontology results for 11,518 region-dependent probes.

**Table S9**. Overlap of differentially methylated probes (DMPs) from cross-region and region-dependent models with AD genetic risk loci.

**Table S10A**. 240 cross-region differentially methylated probes consistent in Lunnon et al. data (p<0.05 and same directionality of effect).[5]

**Table S10B**. 2,908 region-dependent differentially methylated probes consistent in Lunnon et al. data (p<0.025).[5]

**Table S11**. Differential gene expression results for genes with nearby differential methylation, based upon the cross-region model.

**Table S12**. Results for genes for which nearby DNAm associates with gene expression (at nominal significance p<0.05) and the gene is differentially expressed (at nominal significance p<0.05).

### Supplemental Figures

**Figure S1.**
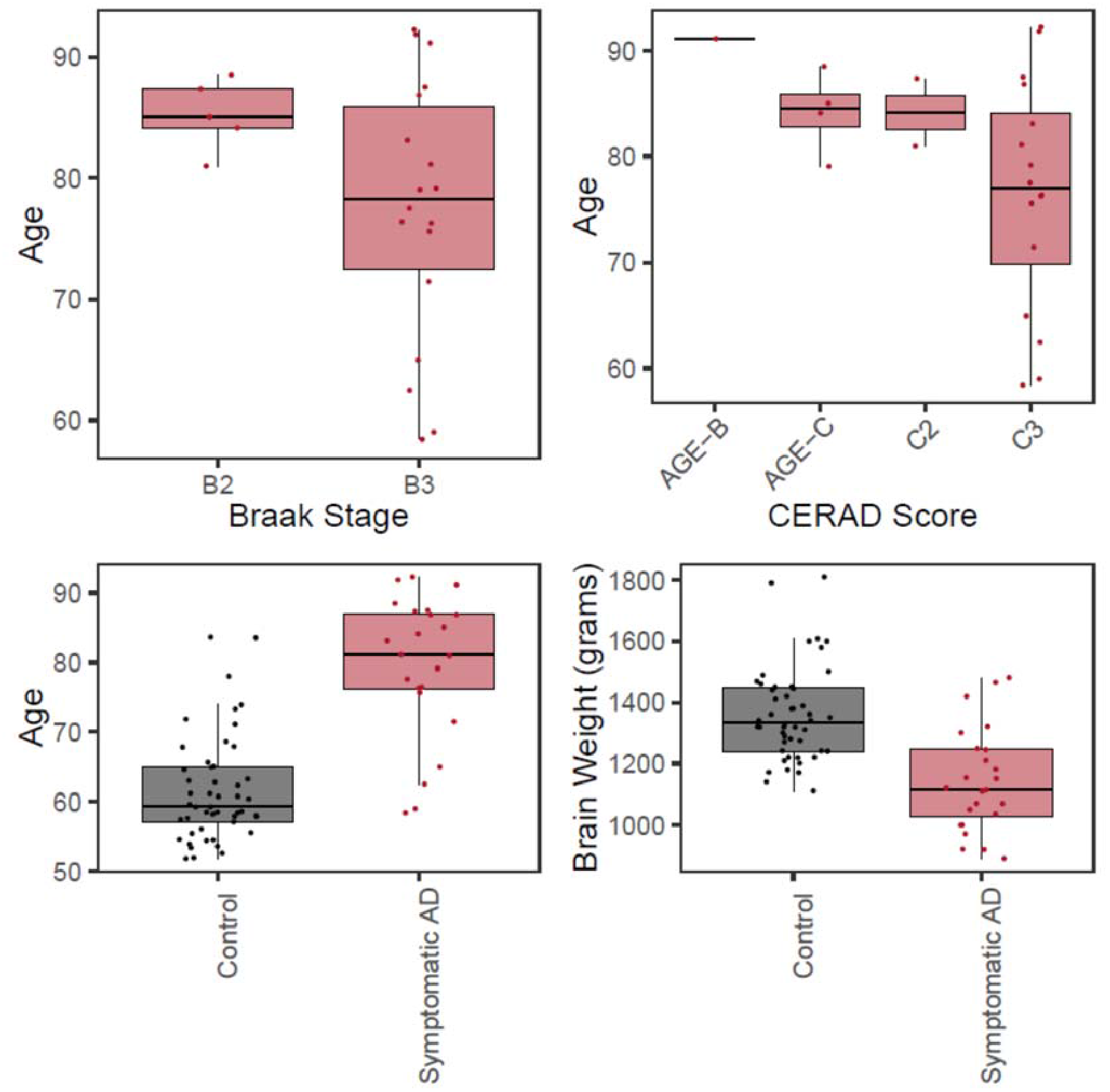
Distribution of demographic variables. Black points indicate control samples and red points/shading indicate symptomatic AD. AD: *Alzheimer’s disease*.

**Figure S2.**
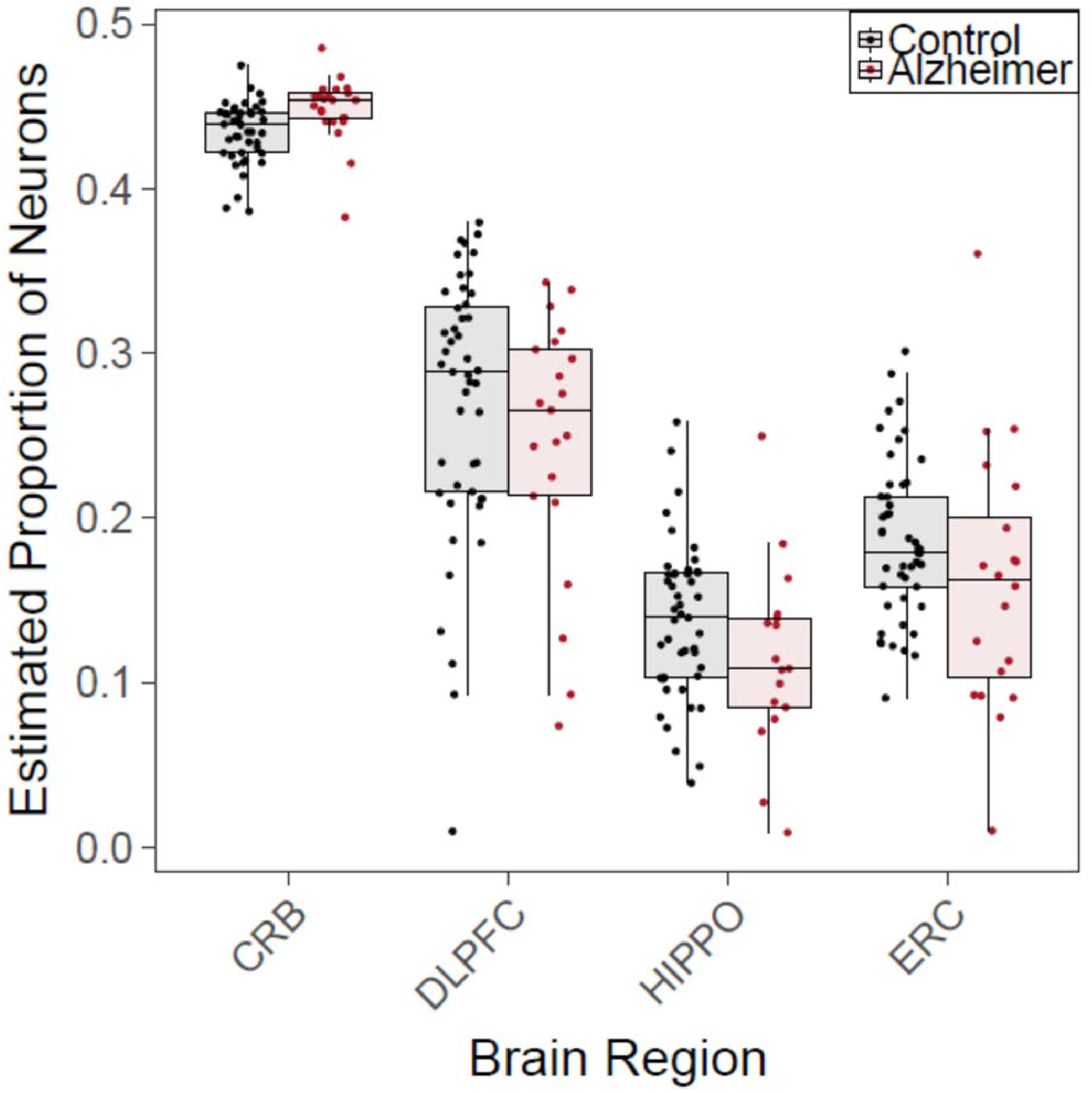
Estimated proportion of neurons in AD and controls across four brain regions. Alzheimer’s disease was not associated with differences in the estimated proportions of neurons in any brain region assessed: CRB (t-statistic=0.29, p=0.77, N_control_=43, N_AD_=24), DLPFC (t-statistic=−0.57, p=0.57, N_control_=47, N_AD_=21), HIPPO (t-statistic=−1.34, p=0.18, N_control_=48, N_AD_=17), and ERC (t-statistic=−0.91, p=0.36, N_control_=49, N_AD_=20; via linear models stratified by brain region and adjusting for age, sex, and ancestry).

**Figure S3A.**
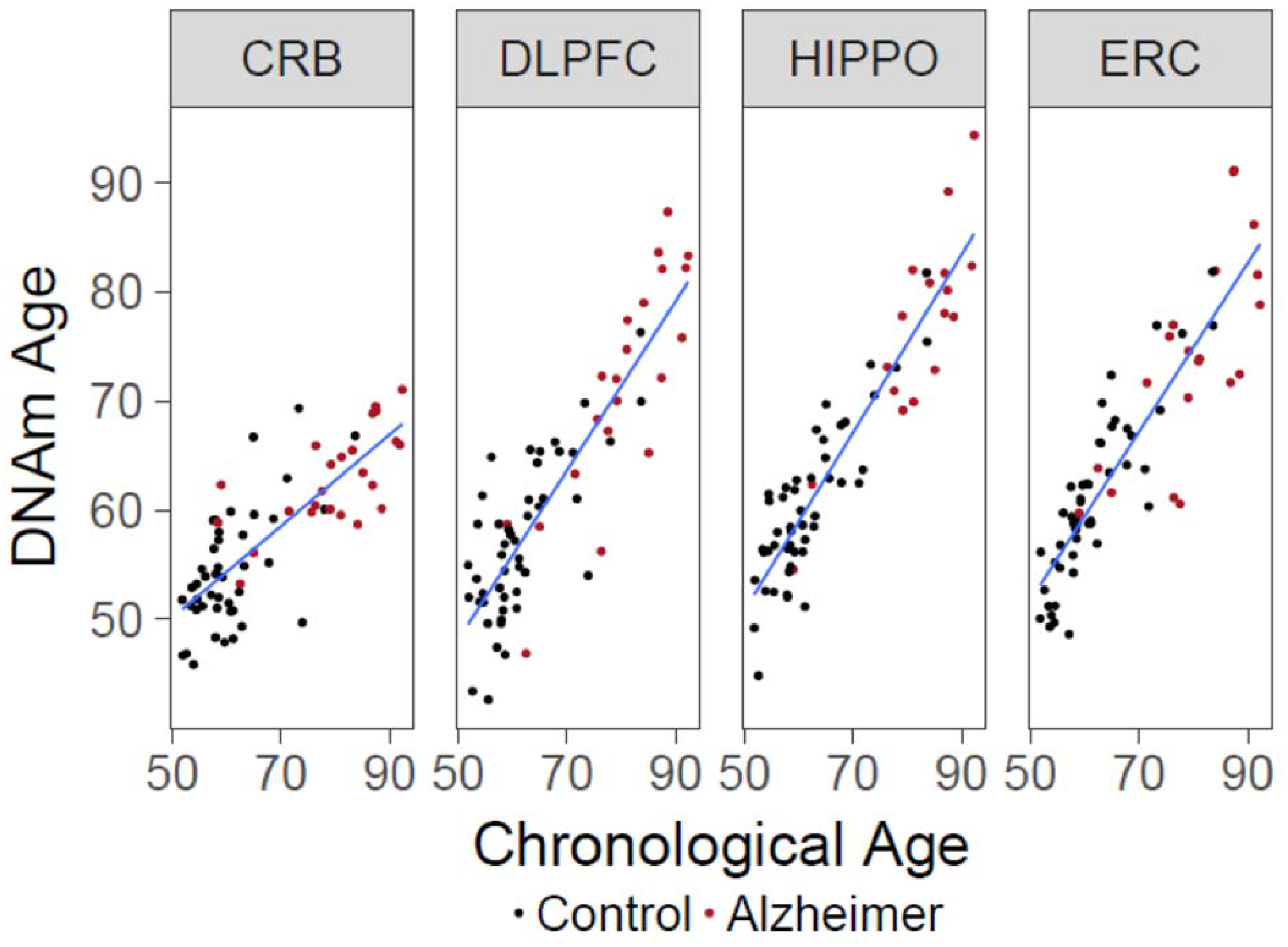
Epigenetic age is highly correlated with chronological age across all four brain regions. Chronological age and DNAm age are highly associated in DLPFC (Pearson’s *r* = 0.871), ERC (*r* = 0.886), and HIPPO (*r* = 0.924, all P<2.2×10^−16^). Cerebellum has a “slower” rate of epigenetic aging as has been proposed previously[19] (*r* = 0.787, P=1.48×10^−16^).

**Figure S3B.**
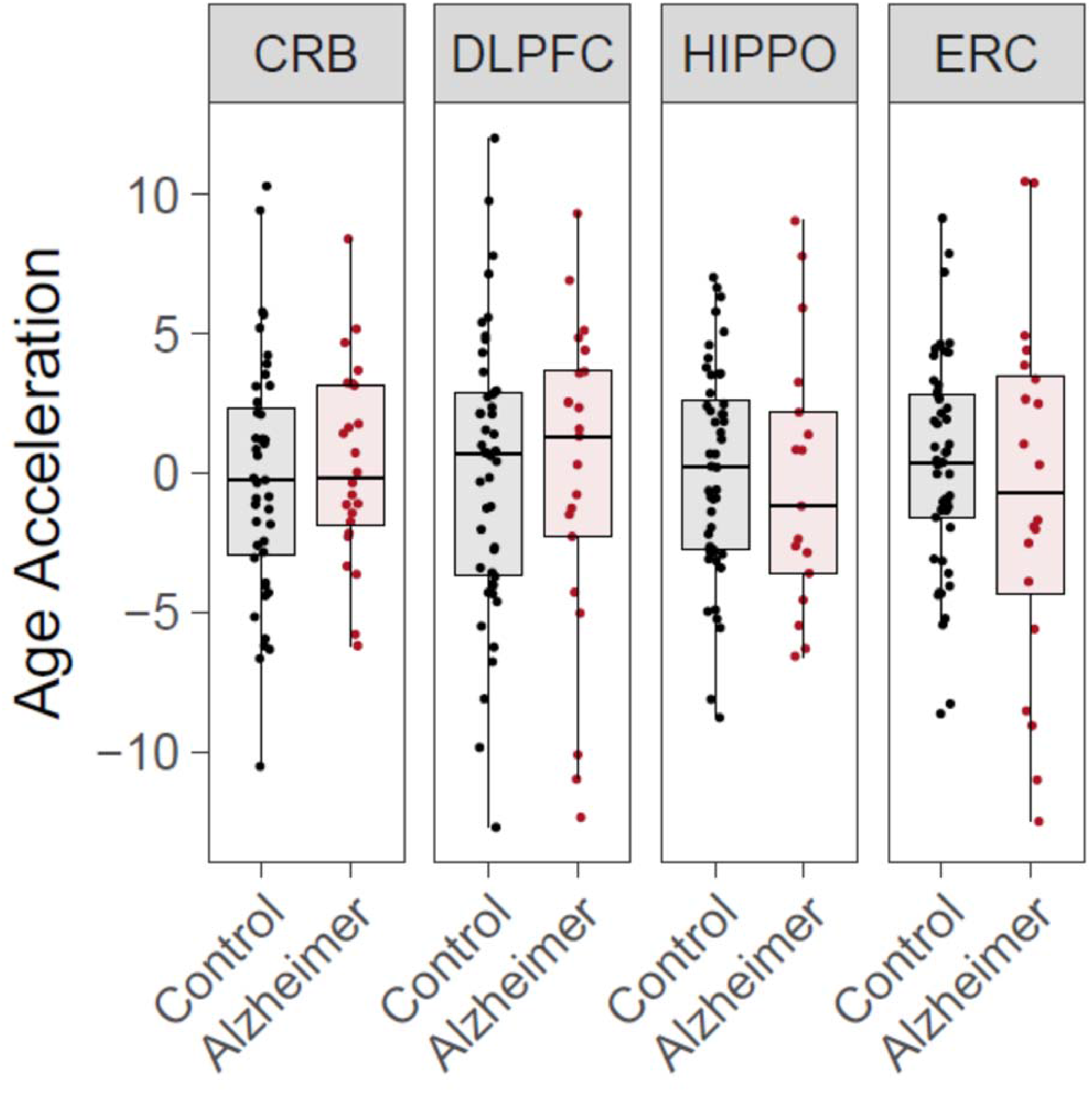
Epigenetic age acceleration and Alzheimer’s disease (AD). We did not find significant differences in epigenetic age acceleration associated with Alzheimer’s disease in any of the four regions we tested (two-tailed t-test): CRB (t = −0.51, p-value = 0.61, N_control_=43, N_AD_=24), DLPFC (t = 0.11, p-value = 0.92, N_control_=47, N_AD_=21), HIPPO (t = 0.25, p-value = 0.81, N_control_=48, N_AD_=17), and ERC (t = 0.66, p-value = 0.52, N_control_=49, N_AD_=20).

**Figure S4 (A-D).**
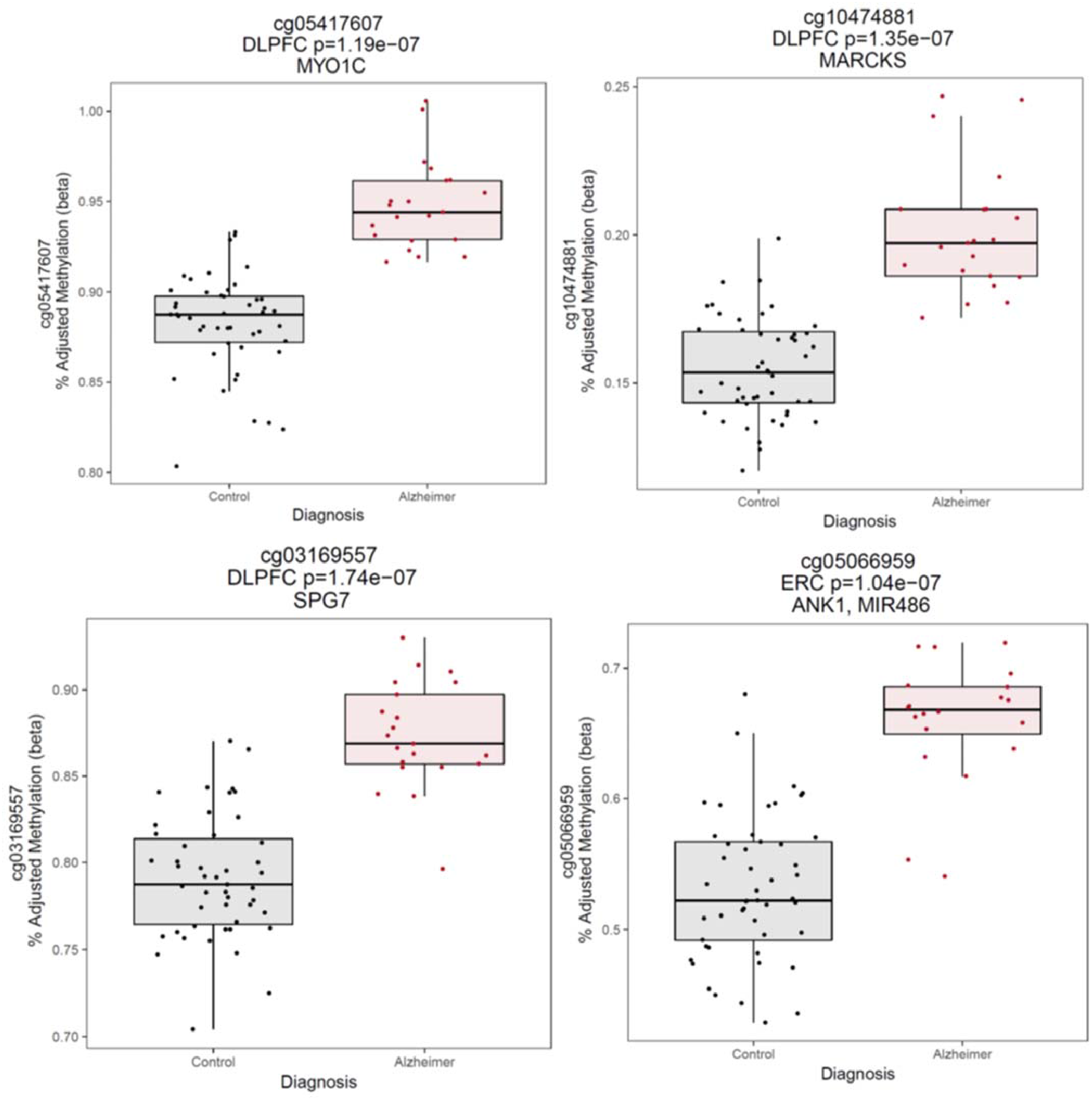
Four differentially methylated probes (DMPs) from the region-specific analysis. Adjusted DNAm values (where the estimated effect of covariates are regressed out) are plotted along the y-axis for both diagnostic groups (x-axis) for four significant DMPs (at false discovery rate, FDR<5%). These four DMPs are annotated to five unique genes *(ANK1* and *MIR486* for cg05066959).

**Supplemental Figure S5.**
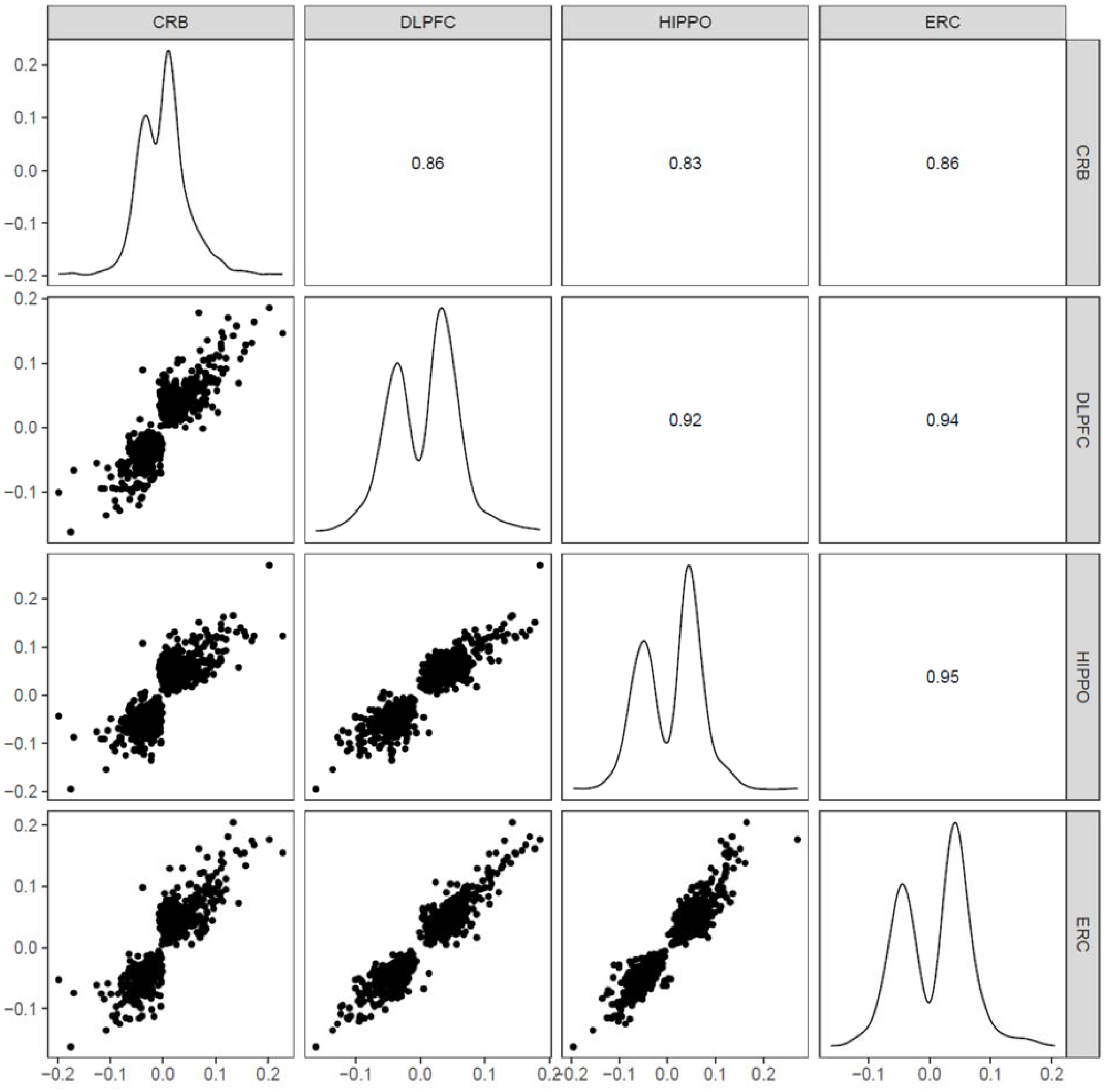
Post-hoc effect sizes of 858 significantly differentially methylated cross-region sites. Pair-wise scatterplots of effect sizes are shown in the bottom right triangular of the plot array. Distributions of the effect sizes are shown along the diagonal. Pearson’s correlations between effect sizes for each brain region are shown in the upper triangular of the plot array.

**Figure S6A.**
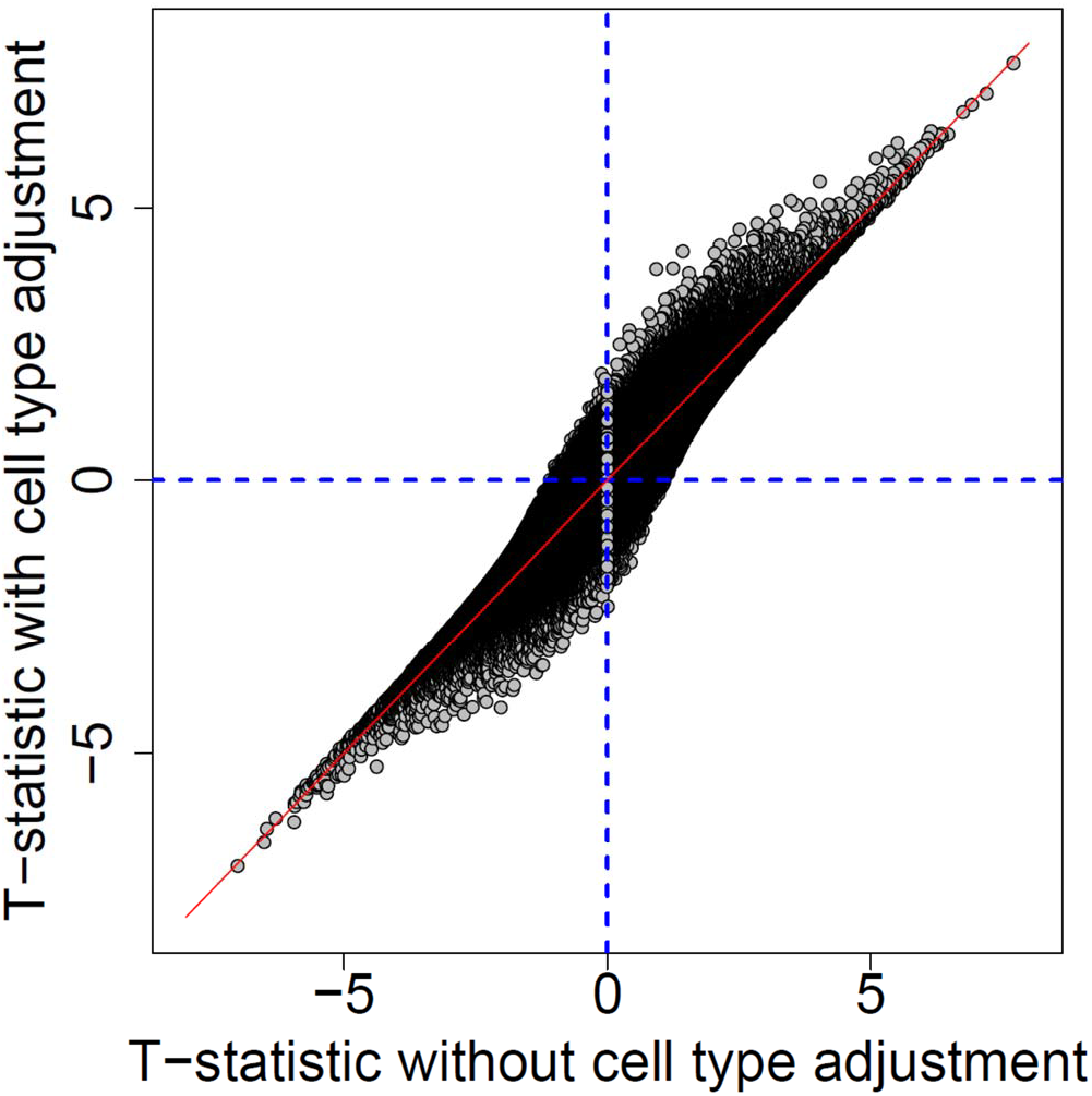
Adjusting for cell-type composition increases t-statistics of AD case-control differences. The solid red line represents the identity line (x=y).

**Figure S6B.**
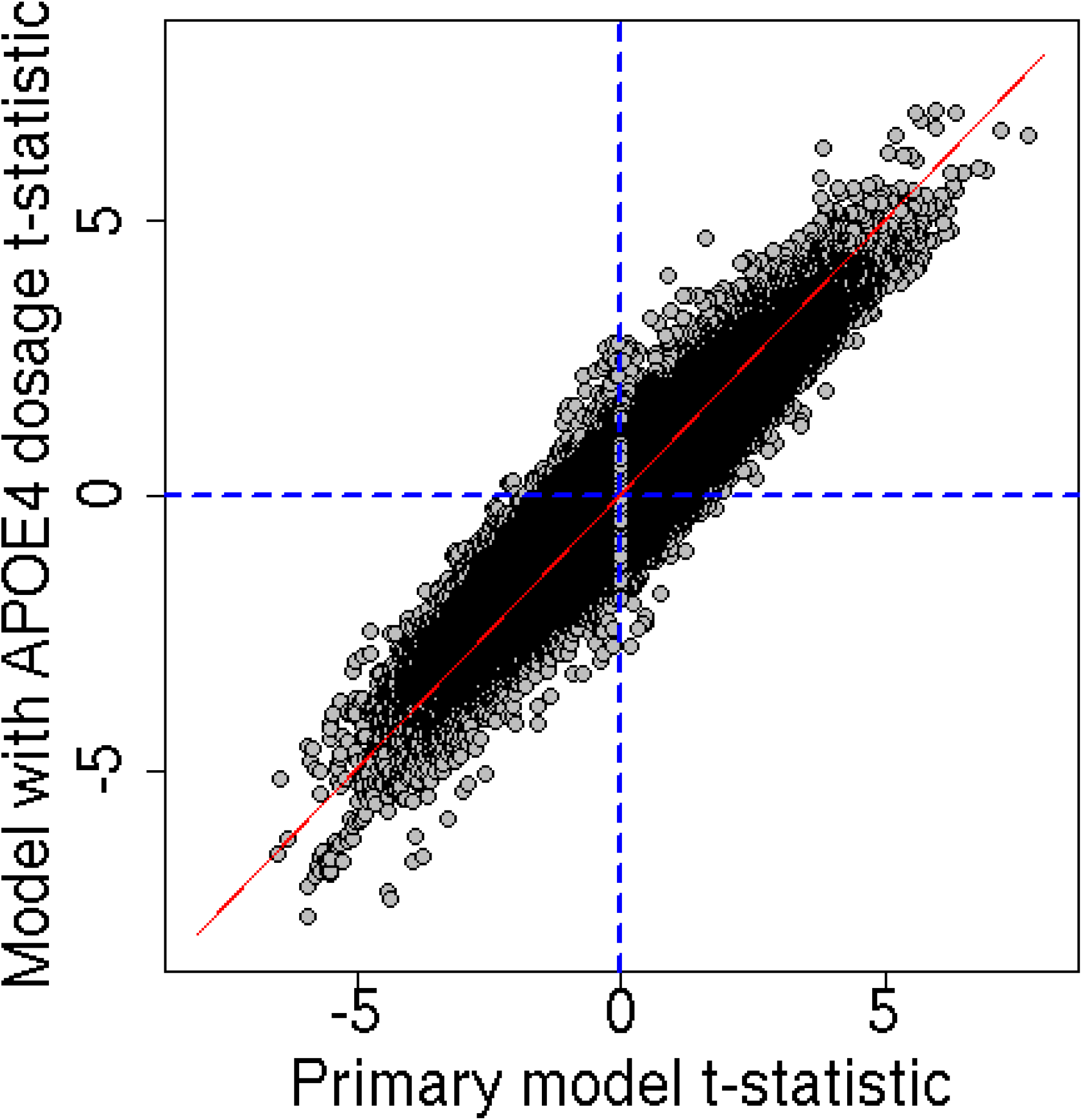
T-statistics from model adjusting for *APOE4* risk burden are highly correlated with the original model.

**Figure S7A.**
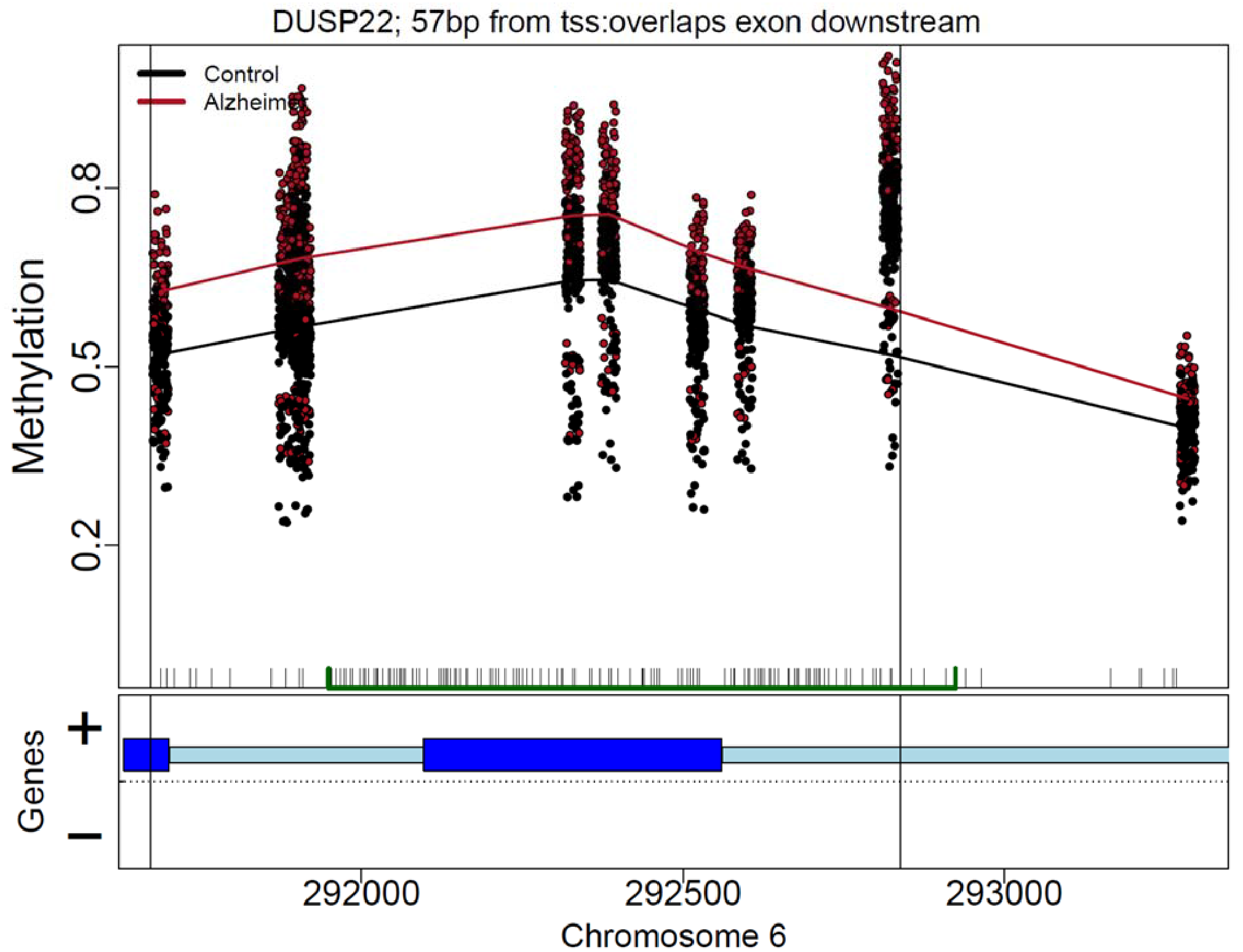
*DUSP22* differentially methylated region (DMR).

**Figure S7B.**
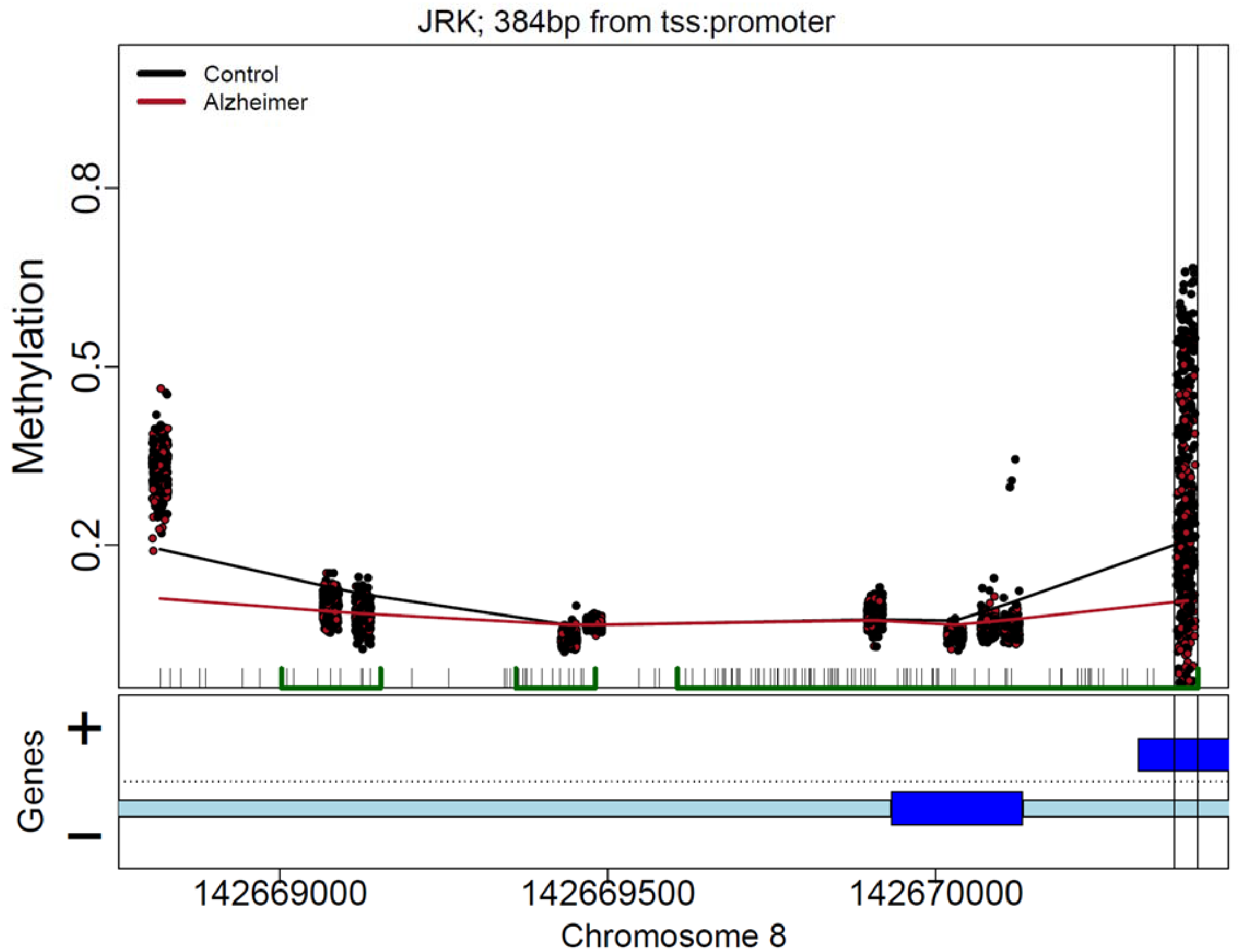
*JRK* differentially methylated region (DMR).

**Figure S7C.**
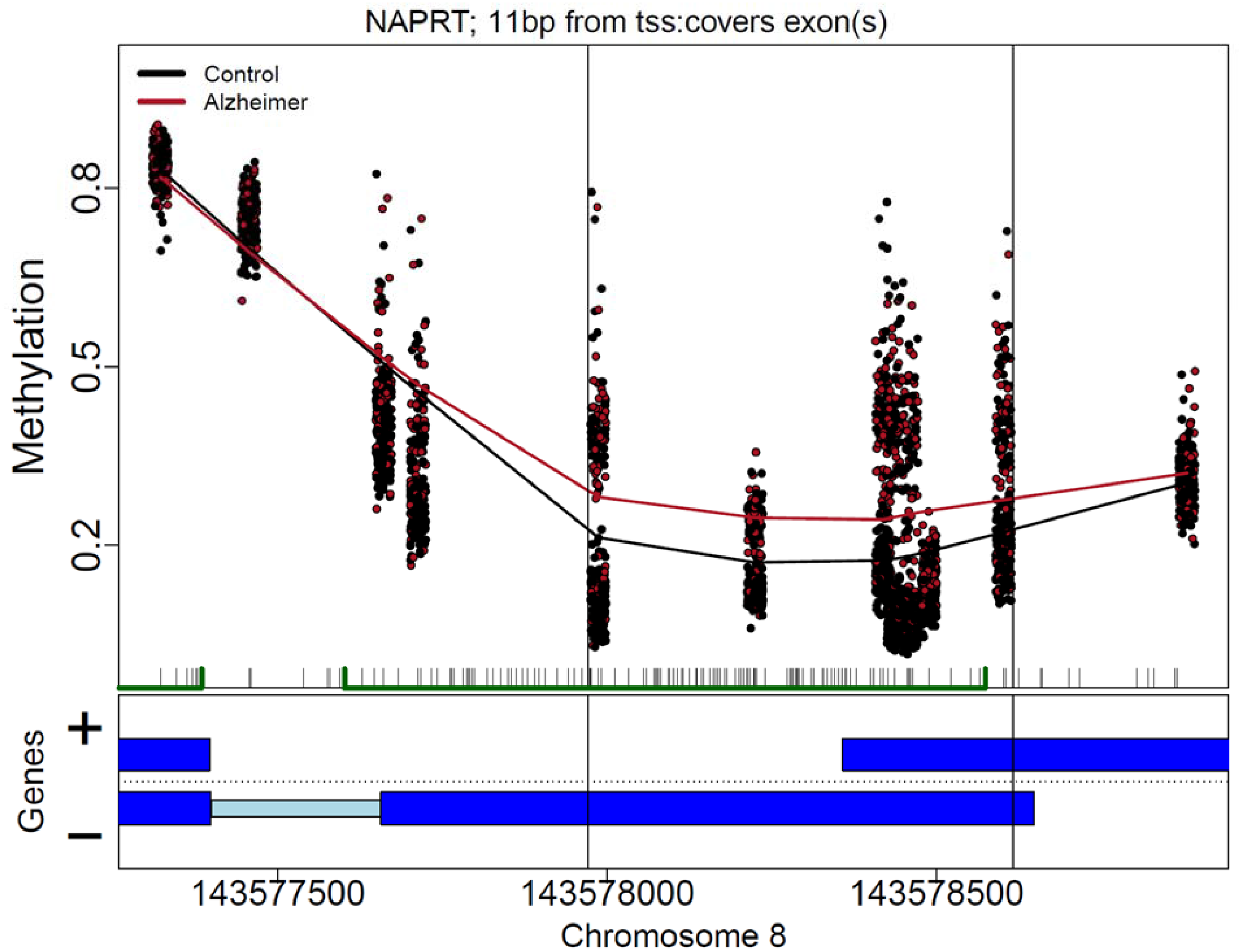
*NAPRT* differentially methylated region (DMR).

**Figure S8.**
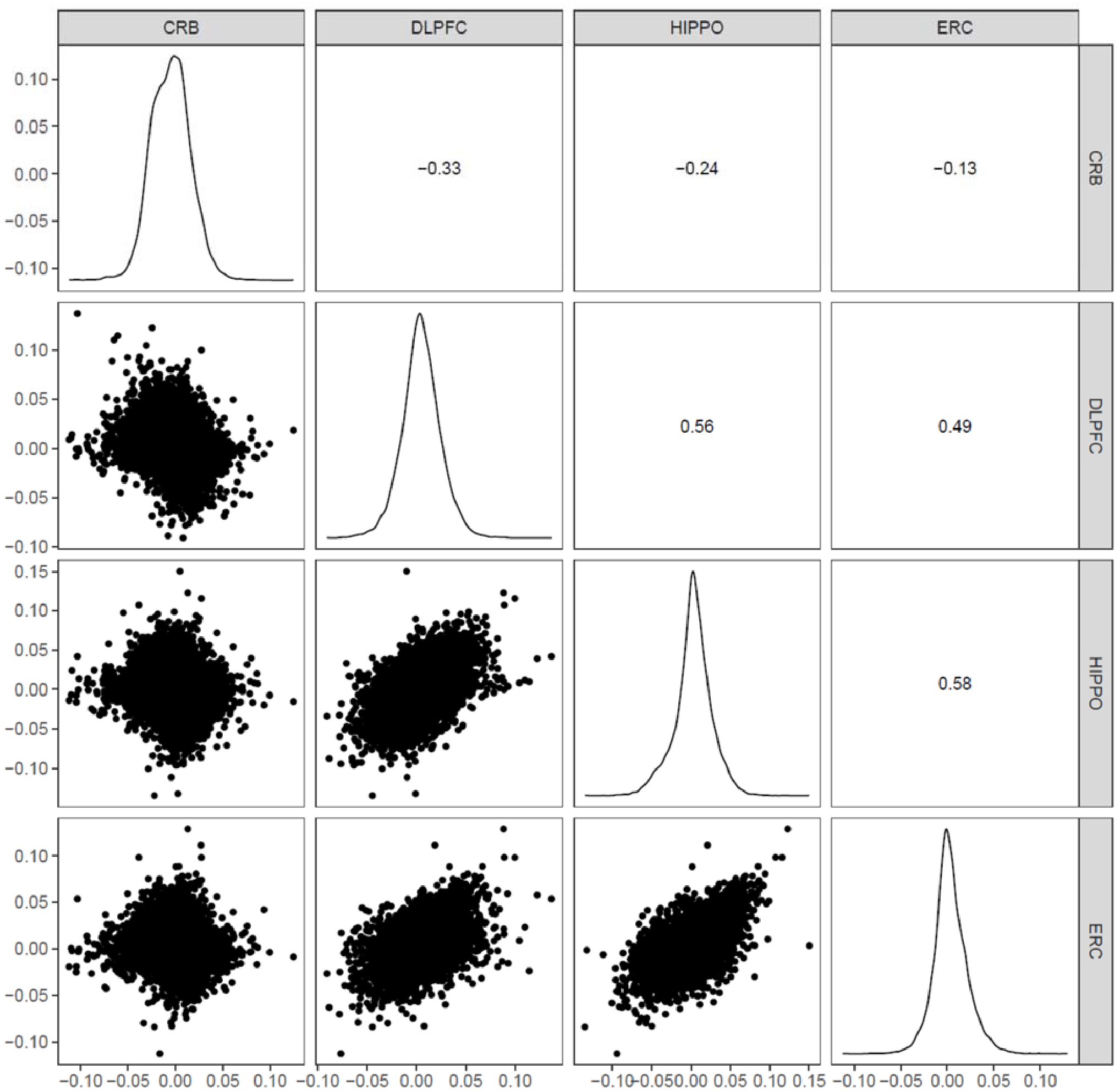
Post-hoc scatter plots of effect sizes from four brain regions for interaction CpGs.

**Figure S9A.**
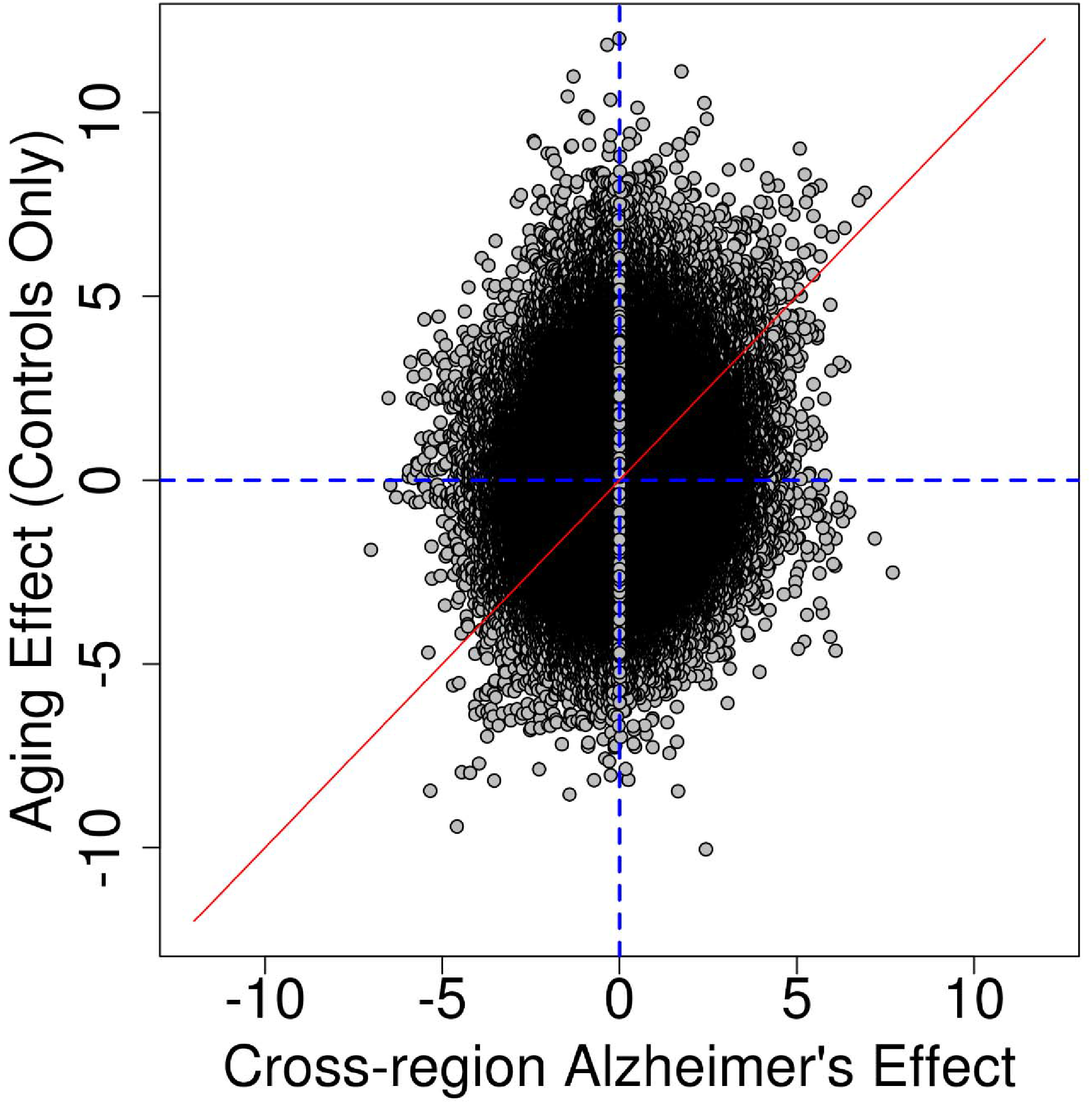
T-statistics from cross-region AD model versus t-statistics from cross-region aging model.

**Figure S9B.**
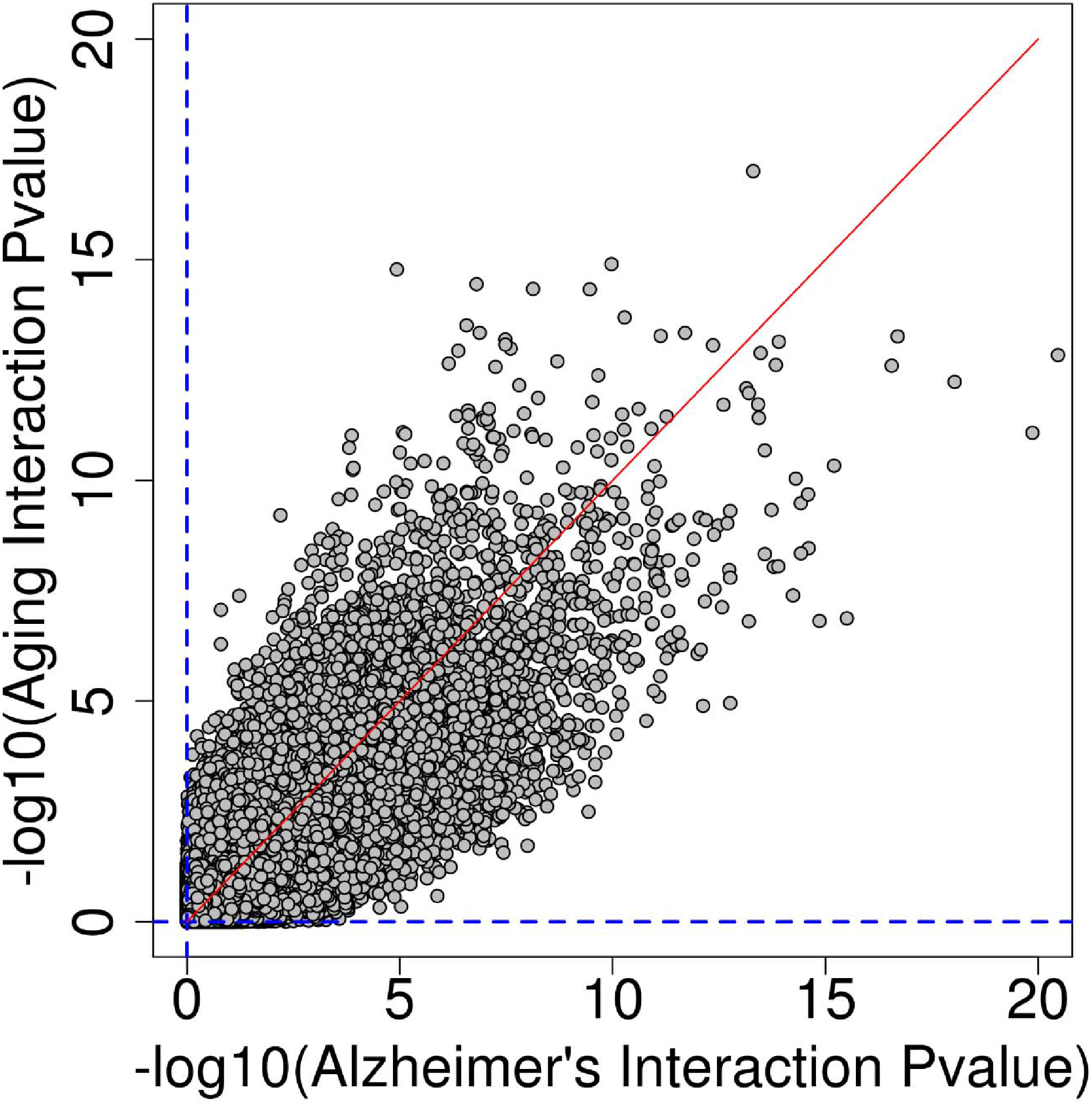
Correlation between −log_10_(Pvalue) from the brain-region by age interaction and − log_10_(Pvalue) from the brain-region by Alzheimer’s disease.

**Figure S10.**
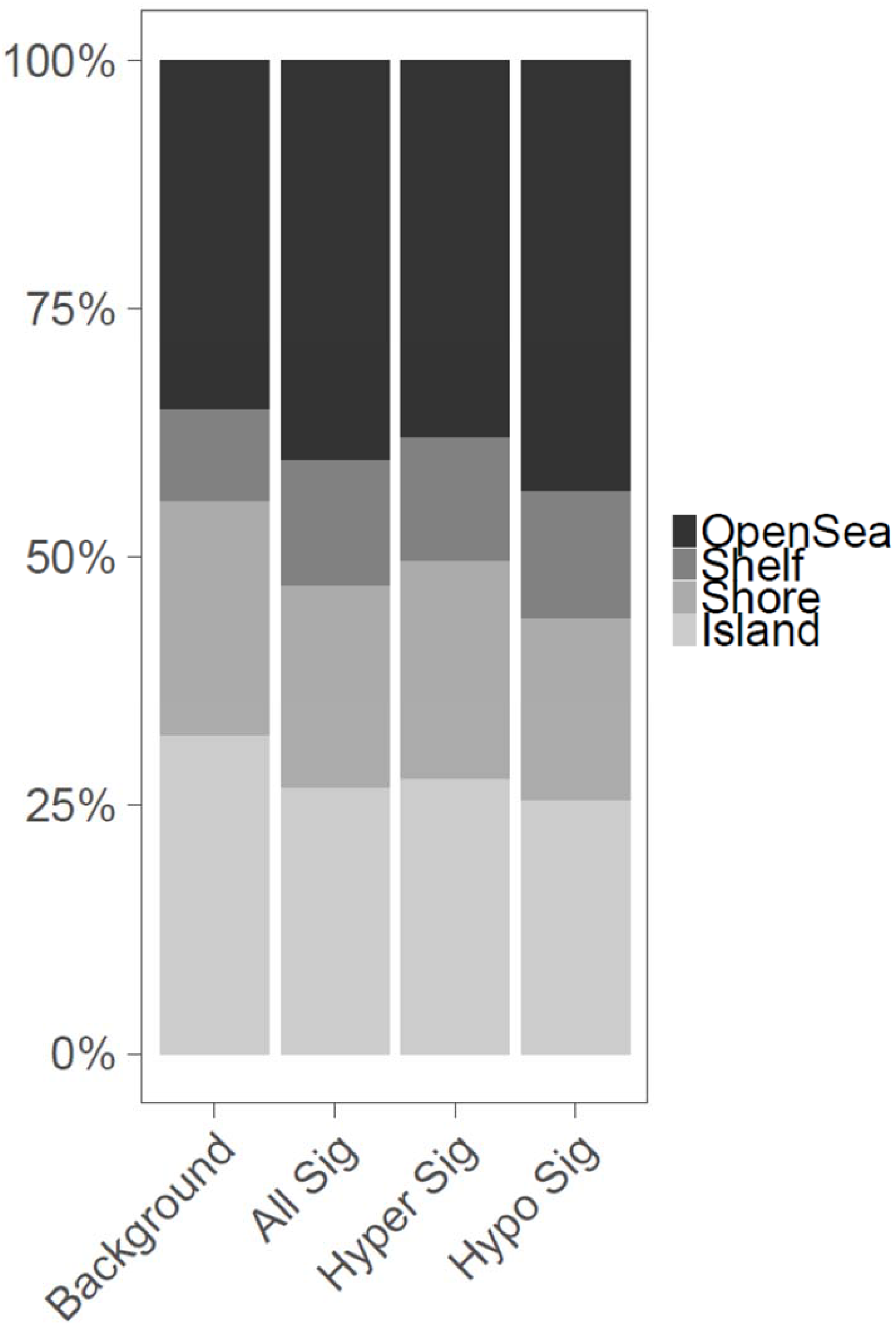
The distribution of CpG genomic density features differed for probes identified as significantly different (FDR<0.05) between AD samples and controls, relative to background (X-squared = 79.23, p<2.2×10^−16^). Overall, in significantly differentially methylated sites, islands were depleted (p=1.55×10^−13^, Odds Ratio, OR=0.645), shores did not differ significantly (p=0.120), shelves were enriched (p=1.35×10^−6^, OR=1.47), and open sea was enriched (p=1.31×10^−7^, OR=1.32).

**Figure S11A.**
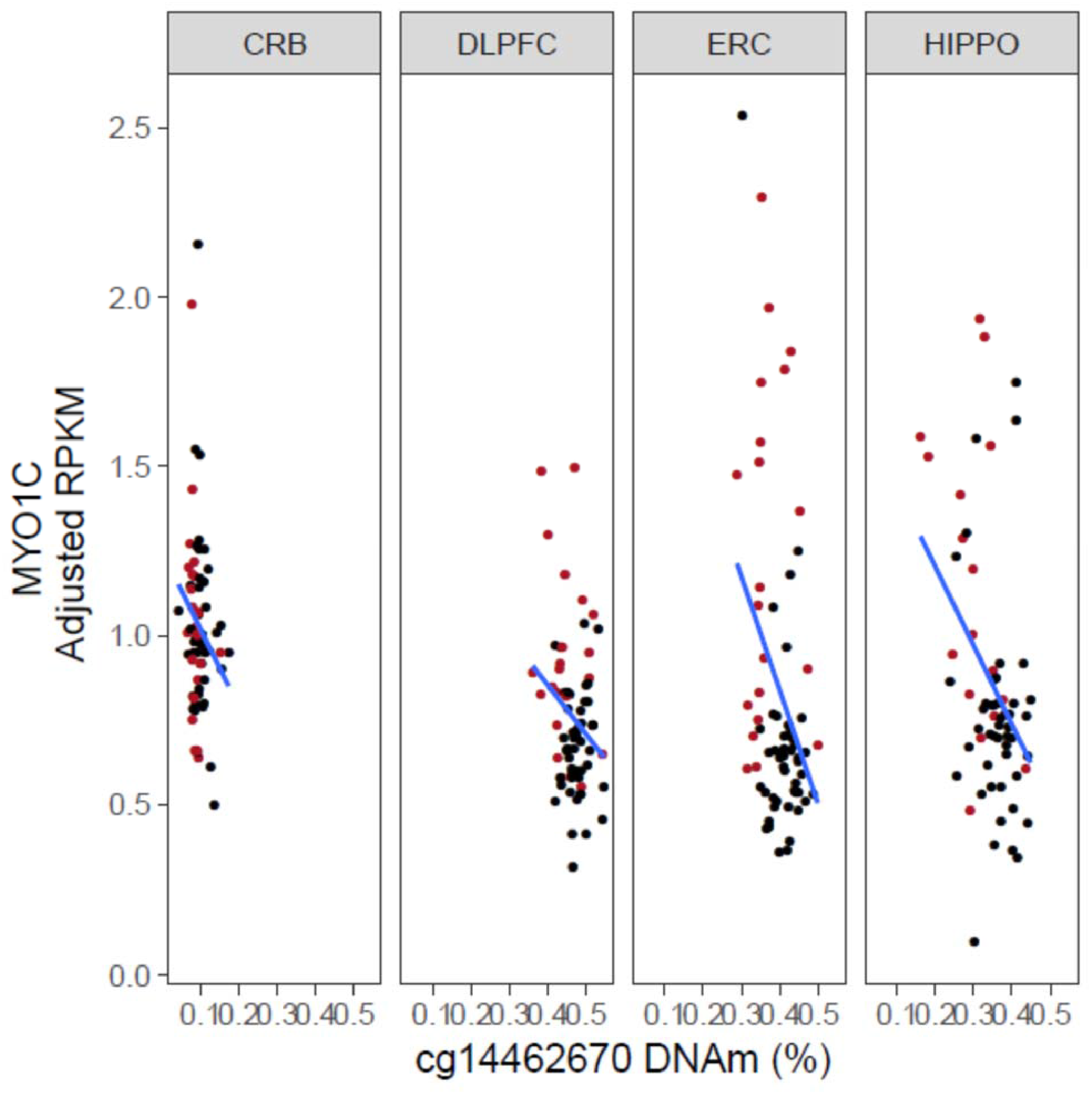
DNAm associates with *MYO1C* expression. Red colored points indicate samples from donors diagnosed with AD; black colored points indicate samples from controls.

**Figure S11B.**
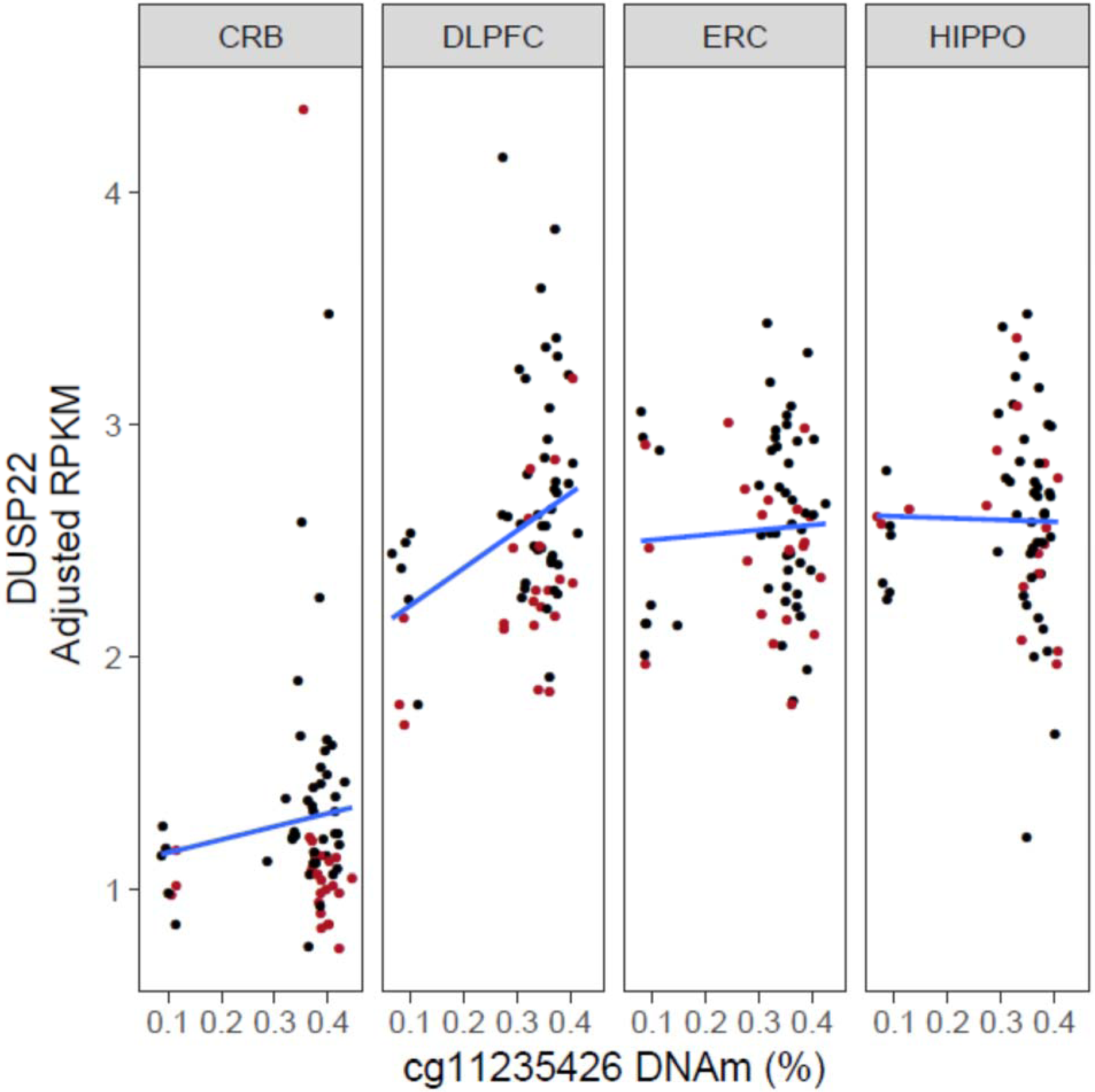
DNAm of cg11235426 associates with *DUSP22* expression only in DLPFC. Red colored points indicate samples from donors diagnosed with AD; black colored points indicate samples from controls.

**Figure S12.**
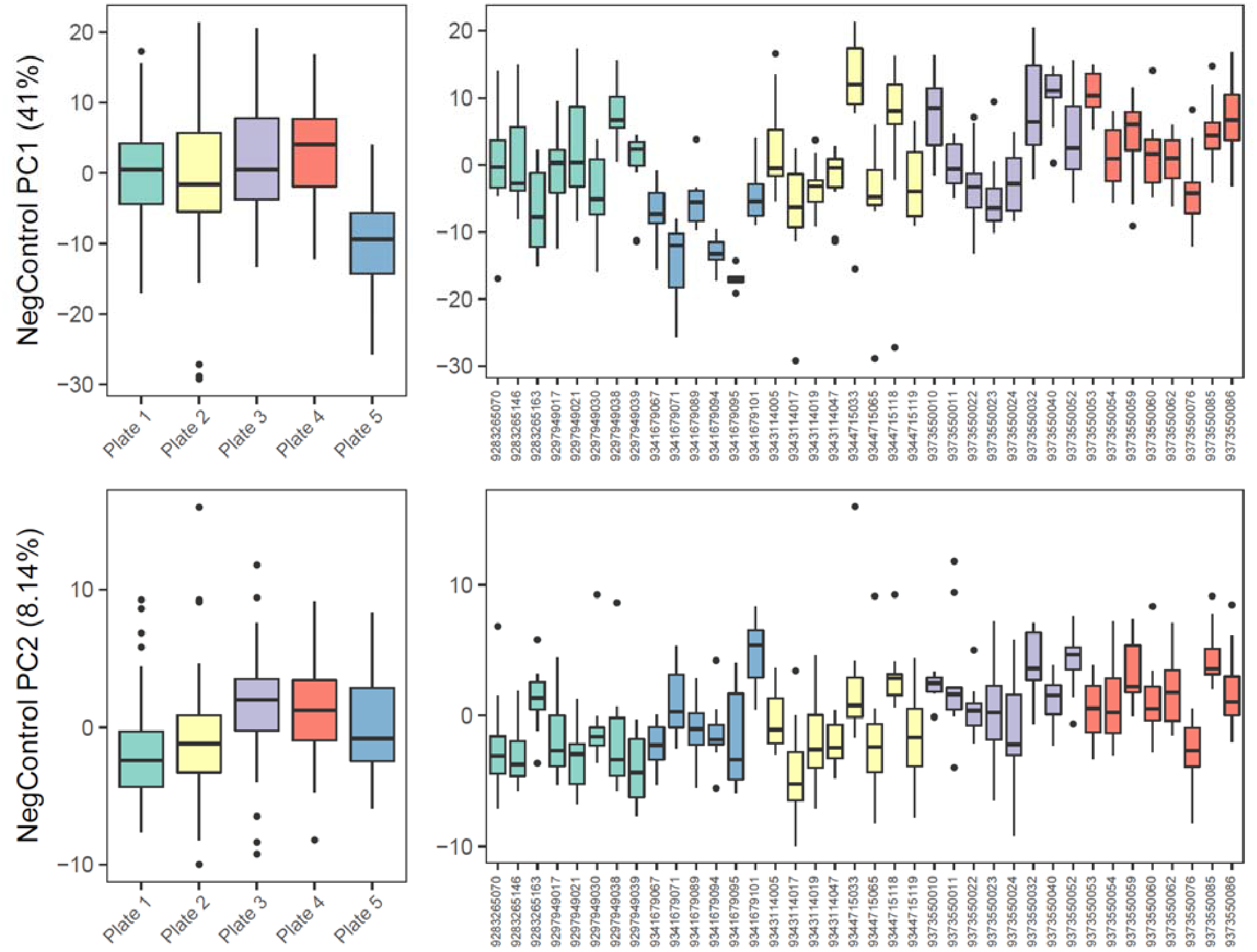
The first two principal components of negative control probe values associate with technical batch effects. Negative control PC1 (41% of variance) associates with sample plate, a technical batch (ANOVA, p<2.2×10^−16^, F-value = 23.92). Negative control PC2 (8.14% of variance) also associates with sample plate (ANOVA, p=1.09×10^−12^, F-value = 16.75).

**Figure S13.**
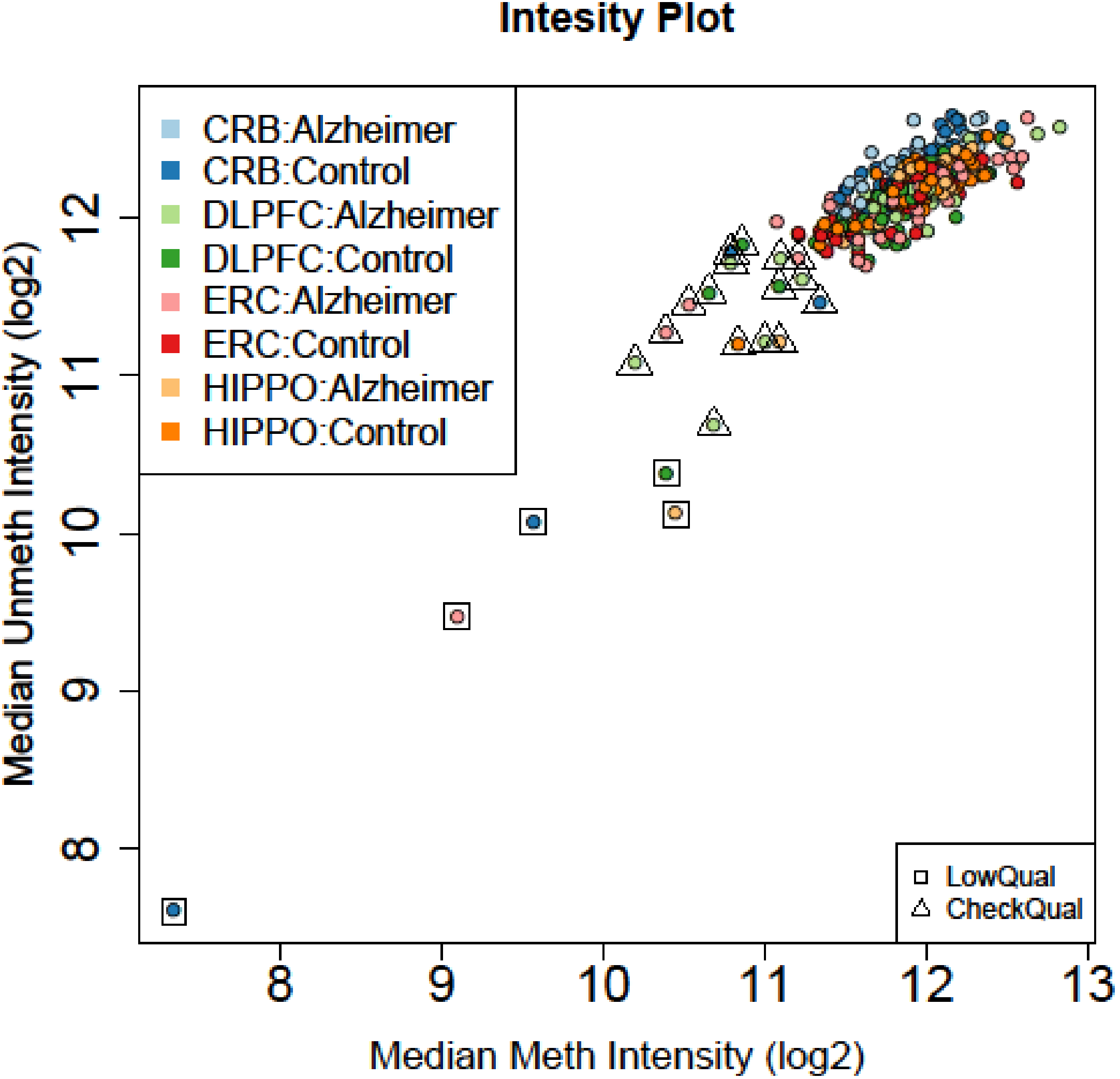
Quality control plot. The median methylation intensity is plotted along the x-axis and the median unmethylation intensity is plotted along the y-axis for all samples (both log2 transformed). Low quality samples (N=5), as indicated by low values of median meth/unmeth, were removed prior to further analysis (boxed and toward bottom left of plot).

**Figure S14.**
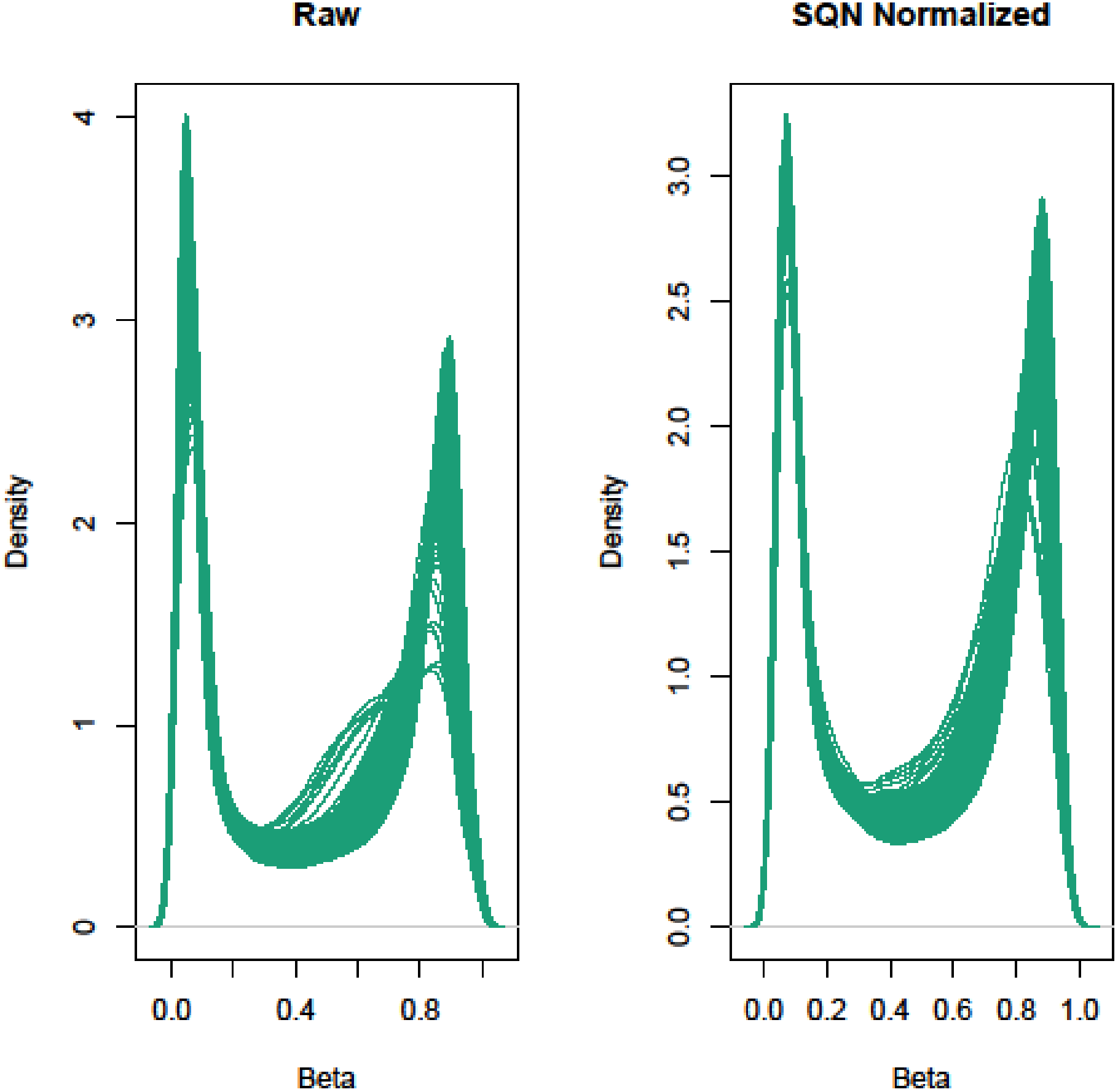
Distribution of beta values before and after stratified quantile normalization (SQN).

**Figure S15.**
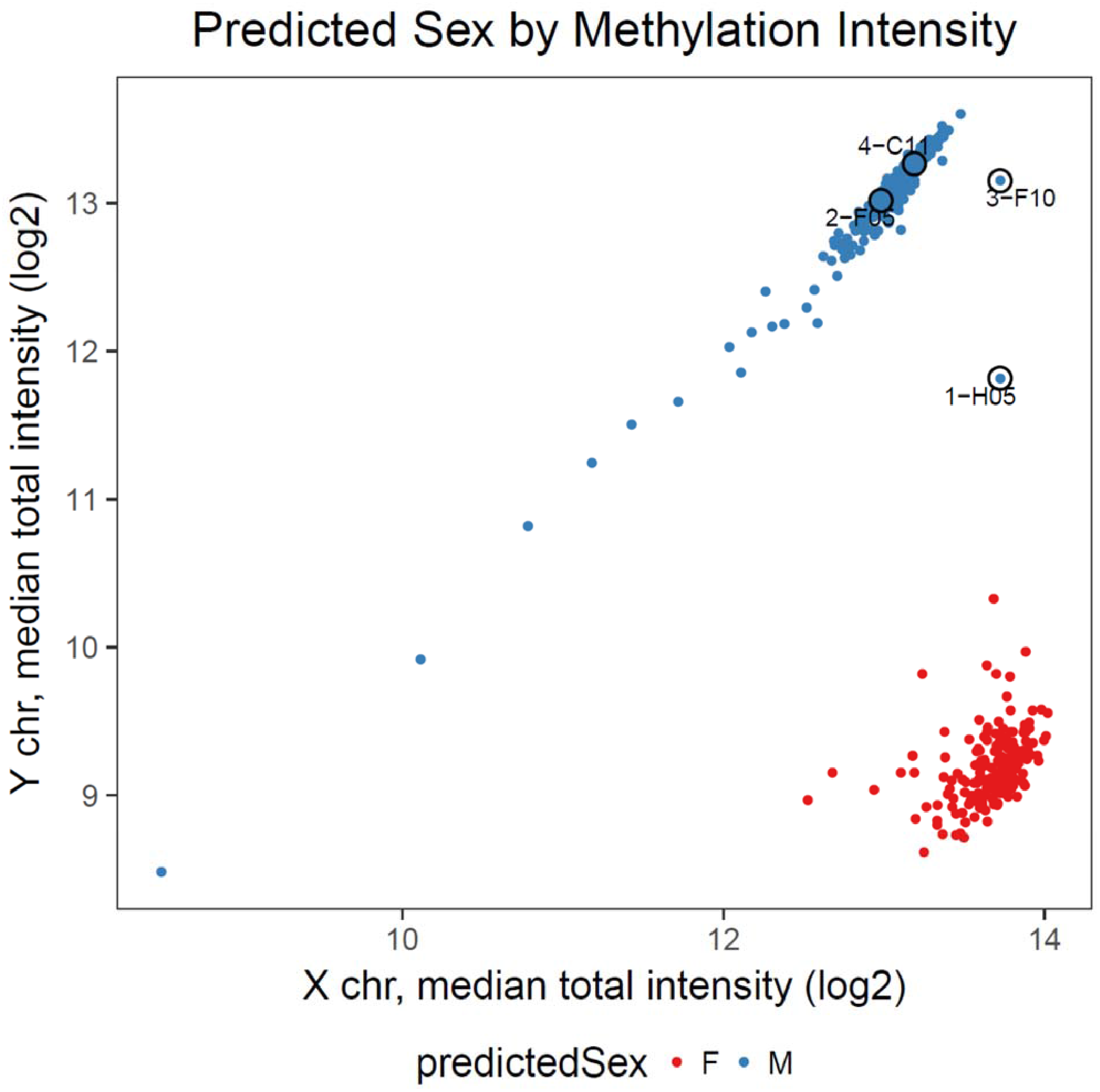
Four potential sex swaps were removed. Four samples (circled and labelled on Figure above) clustered with male samples on the basis of X and Y chromosome median total methylation values, but had a phenotypically reported sex of female. These samples were removed prior to further analysis (N=4).

**Figure S16.**
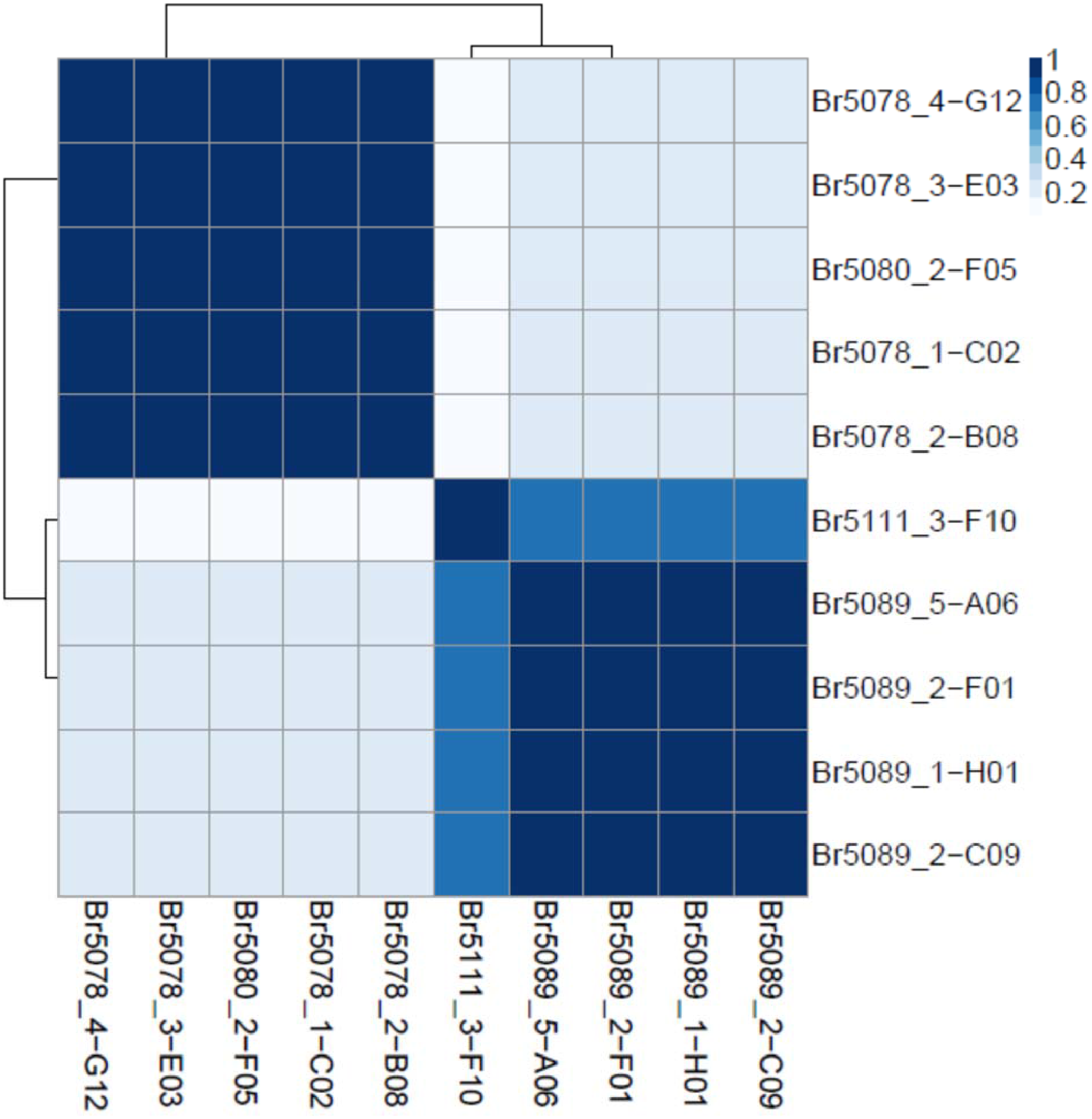
Inappropriately clustering SNP probes on HM450k. Two samples (Br5080_2-F05 and Br5111_3-F10) clustered incorrectly based on genotype and were removed prior to further analysis.

**Figure S17.**
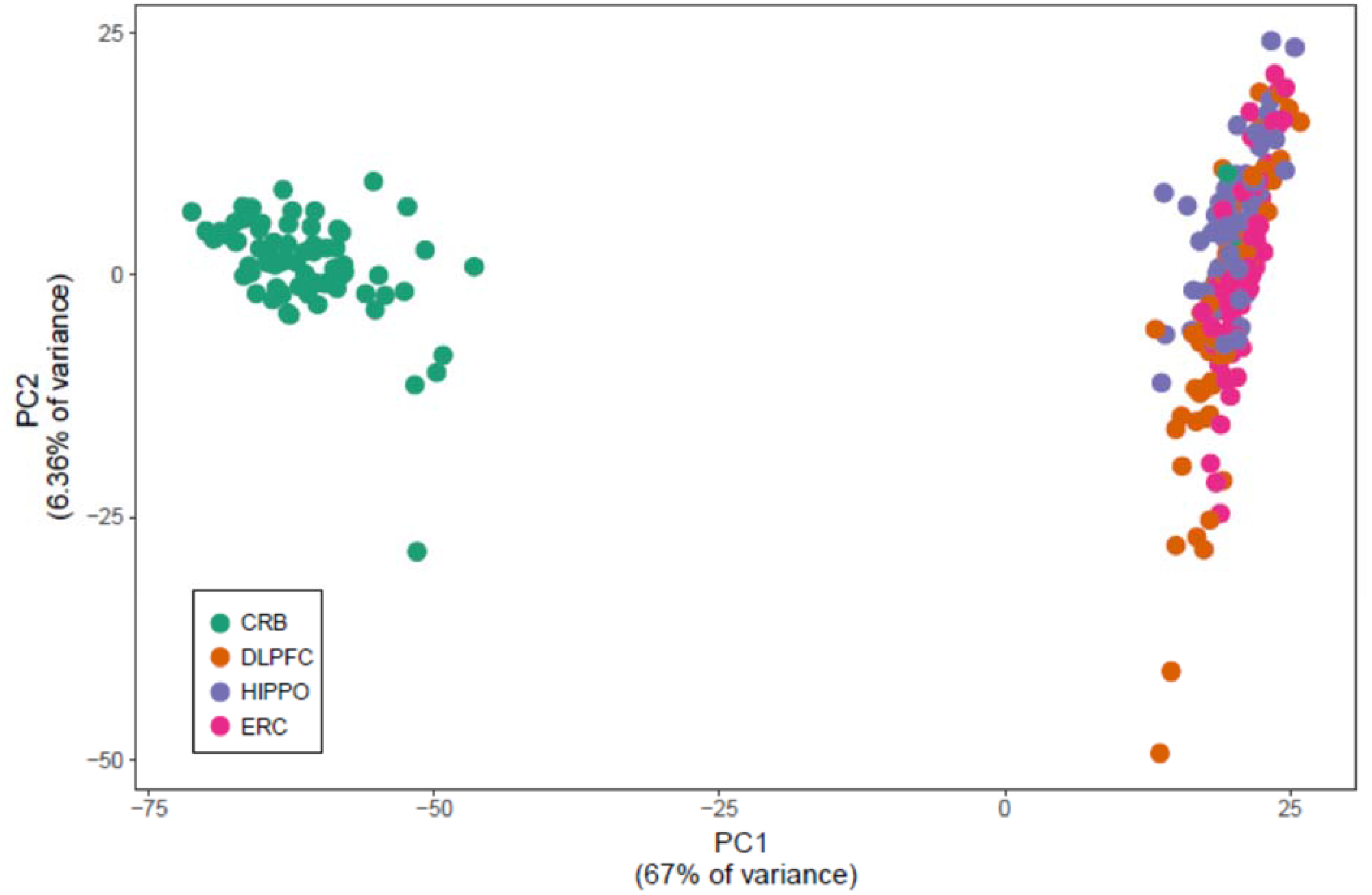
Principal component analysis (PCA). Three samples were labeled as cerebellum but clustered with cortical brain regions on PC1 (~67% of variance). These samples were removed prior to further analysis.

